# Structural basis for inactivation of PRC2 by G-quadruplex RNA

**DOI:** 10.1101/2023.02.06.527314

**Authors:** Jiarui Song, Anne R. Gooding, Wayne O. Hemphill, Vignesh Kasinath, Thomas R. Cech

## Abstract

The histone methyltransferase PRC2 (Polycomb Repressive Complex 2) silences genes via successively attaching three methyl groups to lysine 27 of histone H3. PRC2 associates with numerous pre-mRNA and lncRNA transcripts with a binding preference for G-quadruplex RNA. Here, we present a 3.3Å-resolution cryo-EM structure of PRC2 bound to a G-quadruplex RNA. Notably, RNA mediates the dimerization of PRC2 by binding both protomers and inducing a protein interface comprised of two copies of the catalytic subunit EZH2, which limits nucleosome DNA interaction and occludes H3 tail accessibility to the active site. Our results reveal an unexpected mechanism for RNA-mediated inactivation of a chromatin-modifying enzyme. Furthermore, the flexible loop of EZH2 that helps stabilize RNA binding also facilitates the handoff between RNA and DNA, an activity implicated in PRC2 regulation by RNA.

**One-Sentence Summary:** Cryo-EM structure of RNA-bound PRC2 dimer elucidates an unexpected mechanism of PRC2 inhibition by RNA.

## Main

Many nuclear proteins that necessarily bind chromatin also bind RNA molecules (*1–3*). The binding of RNA has been suggested to facilitate both positive and negative regulation (e.g., complex recruitment and enzymatic inhibition, respectively). PRC2 is a prominent example of a chromatin modifier known to be regulated by RNA (*4, 5*). PRC2 is essential for embryonic development and cell differentiation (*6, 7*). Some tumors are PRC2-dependent (e.g., because of silencing of tumor suppressor genes), making PRC2 a target for cancer therapeutics (*8*). PRC2 consists of four core protein components: EZH2 (catalytic subunit), EED (binds H3K27me3), SUZ12 (provides a platform) and RBAP48 (*7*). Associating cofactors define two PRC2 subclasses (*9, 10*), of which PRC2.2, containing AEBP2 and JARID2, is the subject of this study.

PRC2 binds numerous pre-mRNA and lncRNA transcripts in vitro and in vivo (*11–13*). This broad RNA recognition can be explained at least in part by PRC2 preferring an RNA G-quadruplex (G4) motif (*14, 15*), which could be ubiquitous in the transcriptome from intramolecular and perhaps even intermolecular assemblies (*16*). Intriguingly, proposed models of RNA regulation of PRC2 remain disparate. First, in the ‘Handoff’ model, PRC2 requires RNA for recruitment and occupancy on a unique subset of targeted chromatin (*17*). Remarkably, the direct handoff from RNA to DNA is an intrinsic property of PRC2, as shown by recent biophysical analyses (*18*). Second, the ‘Eviction’ model suggests that nascent RNA removes PRC2 from actively transcribed chromatin to restrict non-specific activity (*14, 19–21*). Third, in the ‘Inhibitor’ model, RNA and nucleosome binding of PRC2 are mutually exclusive, so RNA serves as a direct competitor to prevent PRC2 action (*14, 20, 22, 23*). Another version of the ‘Inhibitor’ model proposes that RNA exploits a regulatory site on PRC2 to abolish H3K27me3 binding EED, which consequently eliminates allosteric activation of EZH2 (*24*). Therefore, structural details of PRC2-RNA interaction have been needed in the field to provide mechanistic insights and coordinate those models.

Cryogenic electron microscopy (cryo-EM) has provided visualization of both substrate-free and nucleosome-bound PRC2 complexes (*25–31*). However, solving a structure of a PRC2-RNA complex has proved challenging. A streptavidin-biotin affinity EM grid approach has been successfully employed in cryo-EM (*32*), and here we adapt this technique for ribonucleoprotein (RNP) complexes using biotinylated RNA. We find that PRC2 can dimerize following RNA binding with a protein-protein interface composed of EZH2 CXC domains. The structure provides a molecular explanation for how RNA acts as a PRC2 inhibitor, and it suggests a mechanism for RNA facilitation of PRC2 recruitment.

### Structure of G-quadruplex RNA-mediated PRC2 dimer

We prepared a six-subunit PRC2.2 complex with full length EZH2 (isoform 2, UniPort Q15910-2), SUZ12, RBAP48, and EED, embryonic-isoform AEBP2, and truncated JARID2_119-450_ (Fig. 1A). We used a streptavidin-affinity EM grid method with 5’ biotinylated G4 RNAs (Fig. 1B), which selects for RNA-bound PRC2 and protects complexes from the hydrophobic water-air interface. Because this method had not been applied to an RNP complex, we validated it by testing different RNA concentrations with the same excess of PRC2. The number of particles observed by negative staining was proportional to the RNA concentration in most fields (Fig. 1C). Unexpectedly, 2D class averages of RNP complexes from streptavidin-selection revealed a significant increase in particles 150Å to 250Å in diameter with profiles showing two recognizable PRC2 molecules (Fig. 1D). We decided to continue with the 1G4 RNA because it exhibited the same 2D projections as 2G4 and seemed more likely to give a single complex.

**Fig. 1.**
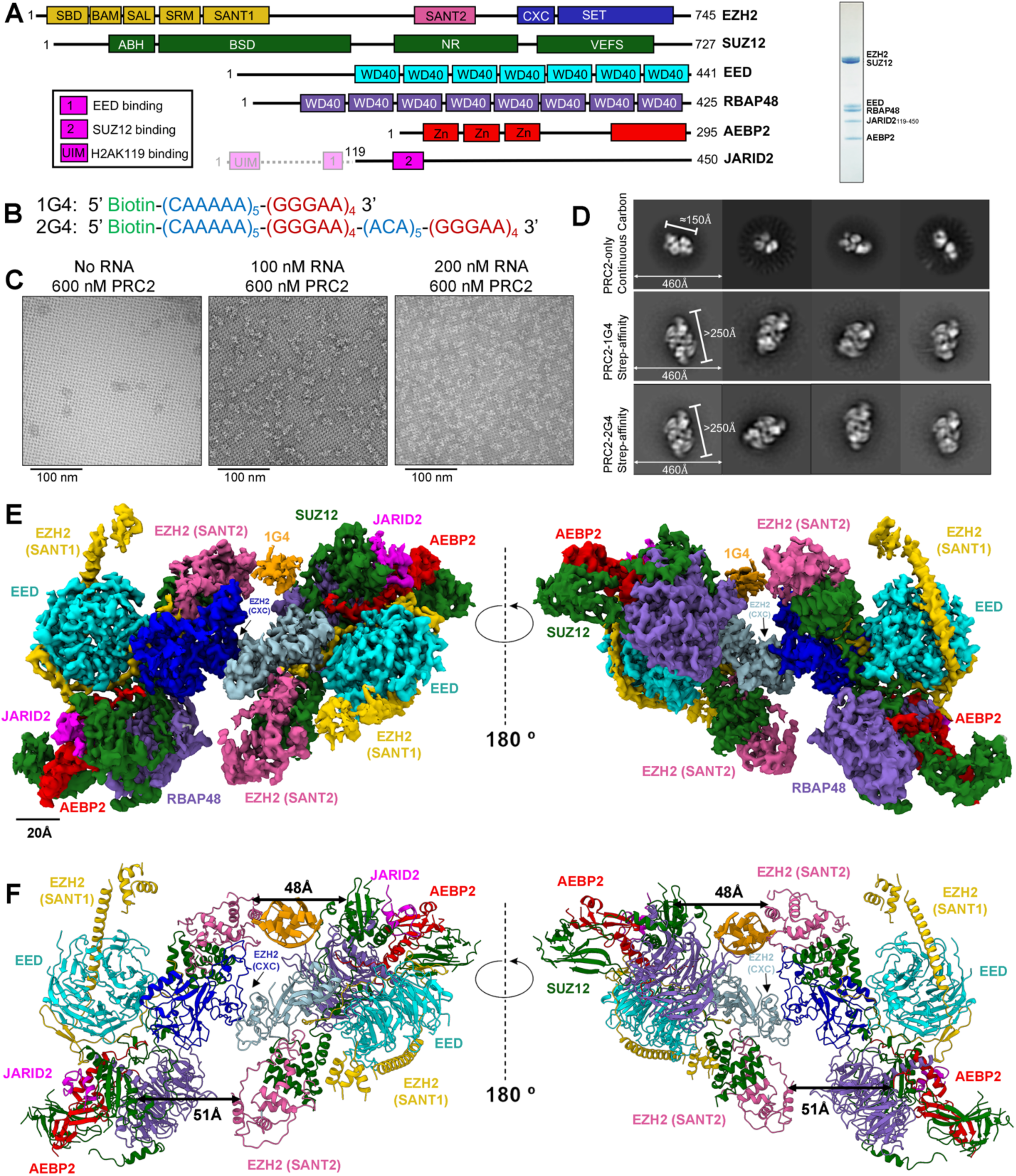
Overall structure of a PRC2-1G4 RNP complex. (**A**) Left: schematic representation of the proteins in the PRC2.2 six-subunit complex. Transparent N-terminal region of JARID2 was not included. Right: Coomassie-stained gel of purified PRC2. (**B**) The two RNA oligonucleotides used in this study. 1G4 has a 5’ biotin, 30-nt (CAAAAA)_5_ A-rich linker sequence, and G4-forming (GGGAA)_4_ sequence. 2G4 has the same 5’ sequence as 1G4 with an additional (GGGAA)_4_ at the 3’ end. (**C**) Negative staining EM images of streptavidin-affinity grids with excess PRC2 and various RNA concentrations. (**D**) 2D-class averages of PRC2 alone collected from continuous carbon grids and PRC2-G4 RNAs from streptavidin-affinity grids. Particle diameters and box sizes are highlighted in white. (**E**) Cryo-EM density map of PRC2 bound to 1G4 RNA. EZH2 (SANT1) is highlighted in gold, EZH2 (SANT2) in hot pink, EZH2 (CXC-SET) of protomer 1 in blue, EZH2 (CXC-SET) of protomer 2 in light blue, EED in cyan, RBAP48 in medium purple, SUZ12 in green, JARID2 in magenta, AEBP2 in red, and 1G4 RNA in orange. (**F**) The atomic model of PRC2 bound to 1G4 RNA. Color scheme is same as (E). Distances between EZH2 (SANT2) and SUZ12 (RRM-like) are highlighted to emphasize the change from 1G4-binding.

We determined the cryo-EM structure of a PRC2-1G4 RNA complex at 3.4Å-resolution from consensus refinement and 3.3Å from multibody refinement (*33*) (Fig. S1 and S2, table S1). The two PRC2 protomers in the RNP complex are nearly identical and have a conformation previously characterized as the SANT1-extended form (*34*) (Fig. 1E, Fig. S3). Notably, the RNA-induced PRC2 dimer has imperfect C_2_ symmetry (Fig. S4A) associated with differential occupancy of RNA in the two symmetric sites (Fig. S4B and S4C). The strong density in one site, attributed to RNA, is fit by a G4 structure (Fig. S4D), but the symmetric site has very weak and discontinuous density which is ignored in the rest of this study. The distance between two PRC2 promoters is reduced from 51Å to 48Å on the side with clear 1G4 density (Fig. 1F), which not only explains the faulty symmetry but strongly supports the model of a single G4 being sufficient for PRC2 dimerization. The additional protein interface in the dimer is a localized EZH2-EZH2 interaction that buries approximately 1740Å^2^ of protein surface (described in subsequent section).

The 1G4 RNA is not nestled into the surface of the protein, as is typically seen for RNA-protein complexes, but instead appears to be separated from the protein. We attribute this to flexible connections between the protein and RNA. The RNA density is in closest proximity to the EZH2 SANT2 domain and the SUZ12 RRM-like (RNA recognition motif-like) domain (Fig. 1F). Those two possible binding sites are supported by previous in vitro and in vivo studies (*24, 25, 35*). We were able to observe clear density linking regions proximal to PRC2 EZH2 SANT2 domain and 1G4 RNA at lower resolution (8-9 Å).

### G-quadruplex RNA induces PRC2 dimerization in solution

To validate the dependence of PRC2 dimerization on G4 RNA binding in solution, we utilized analytical size-exclusion chromatography. In the absence of RNA, our six-subunit PRC2 complex chromatographed as a monomer (Fig. 2), consistent with an absolute molecular weight of 340 kDa measured by mass photometry (Fig. S5A). Incubating PRC2 with 18 kDa 1G4 RNA or 30 kDa 2G4 RNA led to an extremely large RNP complex of approximately 800 kDa, consistent with a dimer (Fig. 2). Microscale thermophoresis (MST) measurements of PRC2-1G4 binding in a G4-favoring KCl buffer exhibited two distinguishable stages of thermophoretic mobility (Fig. S5B). This biphasic binding curve is typical for two binding events (*36*), suggesting that a higher-affinity binding site on PRC2 was primarily occupied at low PRC2 concentrations (1 PRC2:1 RNA), and at higher PRC2 concentrations, lower-affinity binding of a second PRC2 followed (2 PRC2:1 RNA). To test this interpretation, we performed the same MST assays in a G4-destabilizing LiCl buffer or using a G-rich single-stranded RNA with no G4-forming potential (Fig. S5B). Neither experiment gave a distinct biphasic curve, indicating that PRC2 dimerizes specifically on RNA containing at least one G4 motif.

**Fig. 2.**
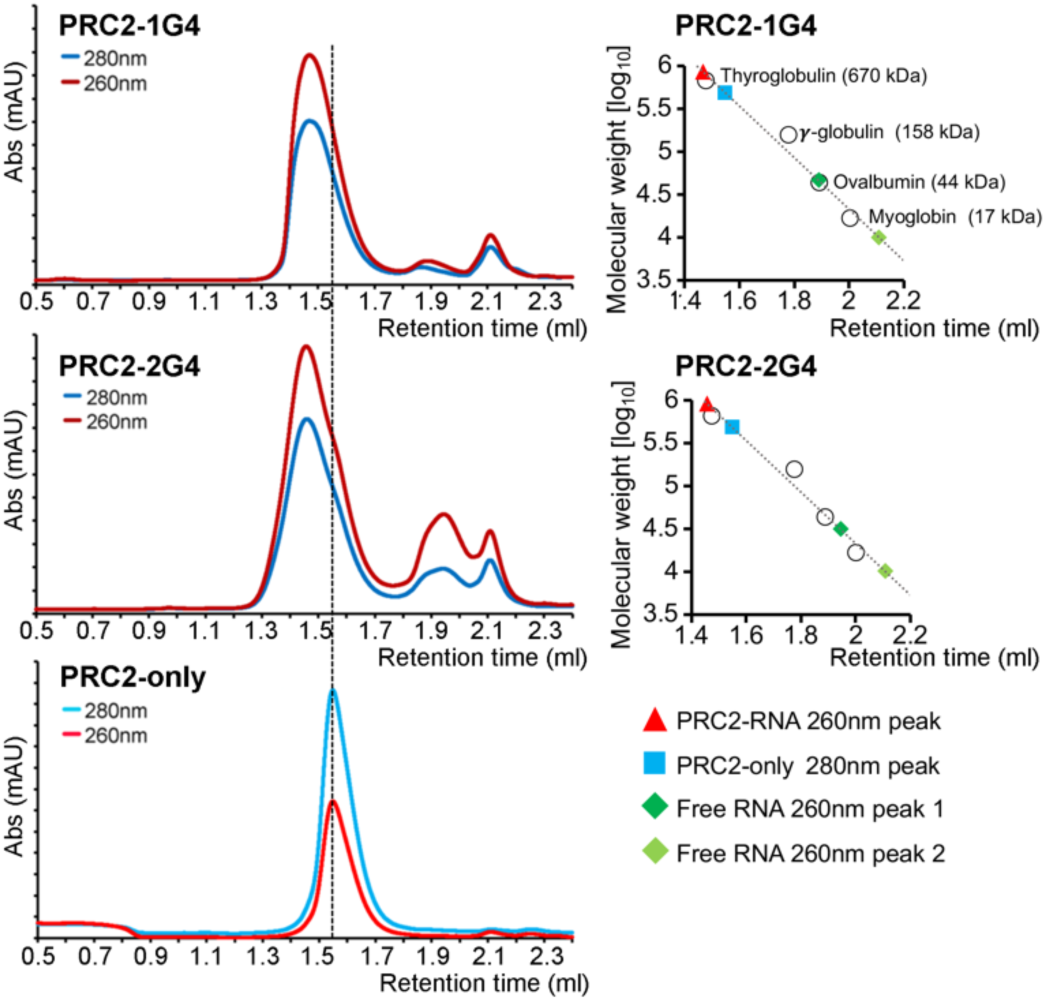
G4 RNA induces PRC2 dimerization in solution. Size-exclusion chromatography of PRC2 preincubated with 1G4, 2G4 and mock. Right: Standard curves were used to estimate the molecular weights of PRC2 complexes.

### The dimer interface prevents nucleosome and H3 tail binding

The PRC2-1G4 RNA structure has three features that would be expected to inhibit nucleosome binding and histone methylation. First, the residues within the CXC domain of EZH2 interact with the CXC of the second protomer to form the dimer interface. This dimer interface includes a K568-K568 hydrophobic interaction, two R566-T573 hydrogen-bond interactions, and two Q575-G564 hydrogen-bond interactions (Fig. 3A). In contrast, in a nucleosome-bound PRC2, the CXC domain facilitates the catalytic activity of the adjacent SET domain of EZH2, specifically with R566 K568 T573 Q575 contributing to interactions with nucleosome DNA and the H3 tail (*26*) (Fig. 3B). The disparate functions of the CXC domain in these different PRC2 structures is seen by superposition of our density map onto the nucleosome-bound PRC2, showing clashes with both the DNA and the H3 tail (Fig. 3C). Therefore, we propose that nucleosome binding and H3 tail-loading – both of which are essential for HMTase activity – are mutually antagonistic with RNA-mediated PRC2 dimerization. The disruption of nucleosome-PRC2 complexes by RNA was confirmed by a competition assay in solution (Fig. 3D).

**Fig. 3.**
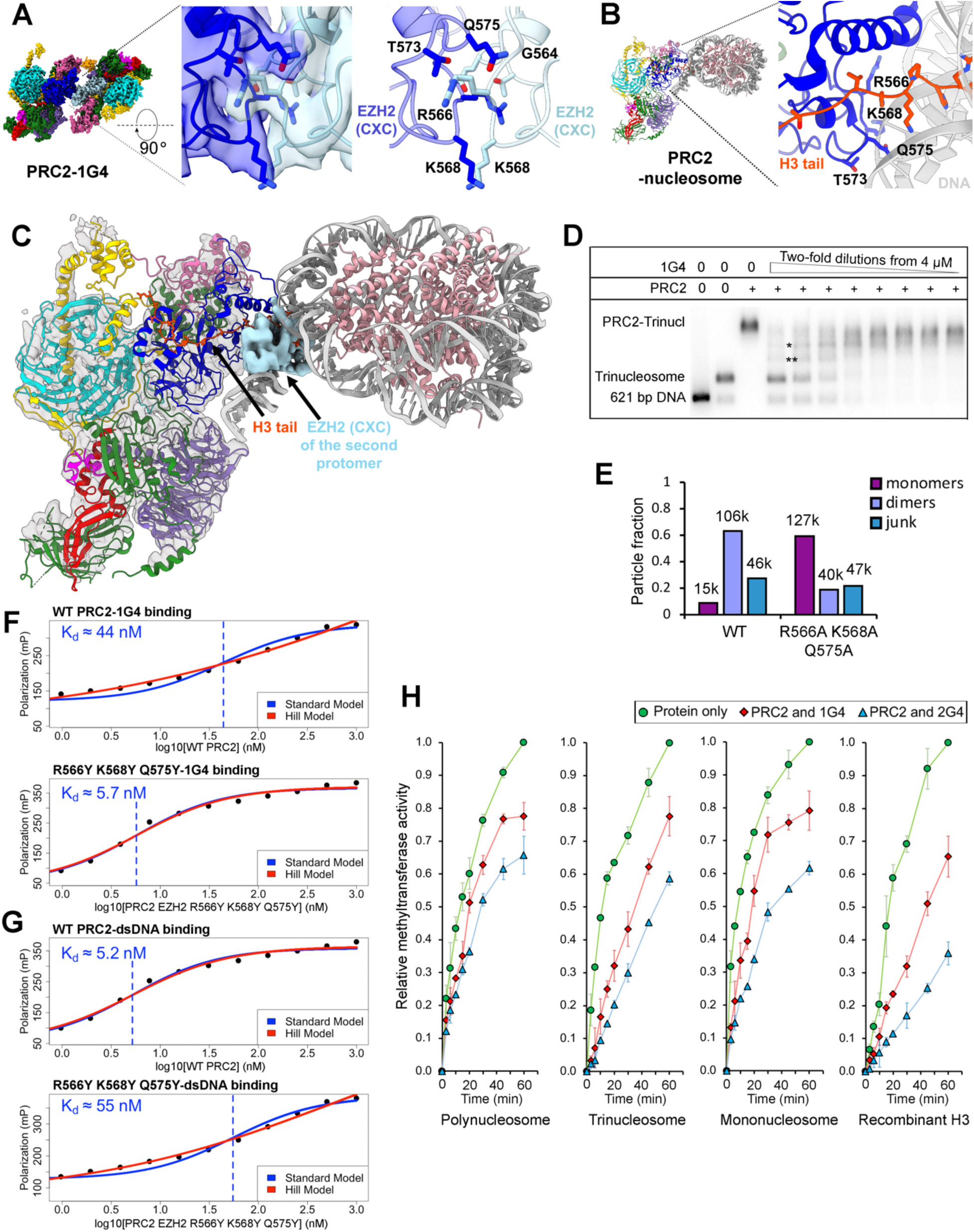
RNA-mediated PRC2 dimer is an inactive complex. (**A**) Cryo-EM structure of PRC2-1G4 complex with zoom-in to show the interface of two EZH2 CXC domains. Interacting residues (R566 K568 T573 Q575) are highlighted in stick representation. (**B**) Structure of PRC2-nucleosome complex (PDB:6WKR) with zoom-in to emphasize the CXC interactions with nucleosome H3 tail. Same residues as shown in (A) are highlighted in stick representation. (**C**) Superposition of the EZH2 CXC domain of the RNA-bound PRC2 on the nucleosome-bound PRC2 to emphasize disparate functions of the CXC domain. (**D**) Nucleosome-RNA competition assay. PRC2 was incubated with constant amount of radiolabeled trinucleosome and serial dilutions of 1G4 RNA. (* and **), incomplete PRC2-trinucleosome complexes. We assume two of three nucleosomes were occupied by PRC2 in * and one of three nucleosomes in **. (**E**) Negative stain EM to quantify monomer and dimer particles of EZH2 R566A K568A Q575A binding 1G4. Number of particles in each class indicated above each bar. (**F**) FP experiments to measure the binding affinity of 1G4 RNA to WT PRC2 and EZH2 R566Y K568Y W575Y. Hill model included the Hill coefficient in the equation to describe cooperativity. (**G**) FP experiments to measure the binding affinity of dsDNA to WT PRC2 and EZH2 R566Y K568Y W575Y. (**H**) Quantification of methyltransferase activities from Fig. S9-S12 are plotted against reaction times for PRC2 (400 nM) preincubated with 1G4 (400 nM), 2G4 (400 nM) and mock (protein only). Error bars are standard deviations of three replicates.

Second, the EZH2 bridge helix (residues 500-516), important for nucleosome DNA binding and channeling H3 tail into the active site of the EZH2 SET domain (*26*), is disordered in both protomers of our RNP complex (Fig. S6). This is consistent with structures of PRC2 lacking nucleosome substrate. Lastly, the EZH2 C-terminal helix (residues 738-742), which points away from the H3K27 binding site, now occludes the active site (Fig. S6) and appears to serve as an additional mechanism to prevent H3 tail binding.

To test the functional importance of the CXC dimer interface, we expressed and purified a mutant (EZH2 R566A K568A Q575A). Mutation of those residues did not affect G4 RNA binding (Fig. S7A), as expected because they do not directly interact with the RNA. However, negative staining EM of streptavidin-affinity grids classified substantially more monomer-size particles from the mutant (59%) than the wild-type (WT) PRC2 (9%) (Fig. 3E, Fig. S7C). This suggests that these EZH2 mutations impacted the overall stability of the PRC2 dimer by destabilizing the protein-protein interaction, and consequently, one protomer dissociated during stringent washes in our grid preparations. Because streptavidin-affinity grids only retain RNA-bound complexes, those monomer particles of the mutant could also represent an intermediate stage of one PRC2 engaging RNA prior to association of the second PRC2 driven by CXC-CXC interaction.

We next attempted to disrupt the CXC interface more severely by substituting bulky sidechains of tyrosine, so we constructed an EZH2 R566Y K568Y Q575Y mutant. Surprisingly, this mutant had higher binding affinity to the G4 RNAs, as determined by fluorescence polarization (FP) assays and electrophoretic mobility shift assays (EMSA) (Fig. 3F, Fig. S8). Notably, double-stranded DNA (dsDNA) binding of this mutant was instead compromised (Fig. 3G), further indicating that DNA and RNA utilize separate mechanisms to engage PRC2 even though they bind mutually antagonistically. Although this mutant is a monomer as observed by negative staining EM, the class averages of RNA-bound particles showed a dominant population of dimer complexes, consisted with the increased RNA binding affinity and the role of RNA in mediating PRC2-dimerization (Fig. S8). We propose that the aromatic sidechains of tyrosine might stack on each other and therefore stabilize the dimer interface. Thus, the dimerization interface need not be very specific, and it appears to be RNA binding rather than protein-protein interaction that drives PRC2 dimerization. Overall, we observed a positive correlation between PRC2 dimerization and G4 RNA binding.

### RNA-induced PRC2 dimer is inactive

Structural observations on the EZH2 CXC interface prompted us to hypothesize that the HMTase activity of the G4-induced PRC2 dimer would be inhibited. To test this, we performed activity assays to compare free PRC2 with RNA-sequestered dimers (Fig. 3H, Fig. S9-S12). PRC2 substrates included HEK293-extracted polynucleosomes, home-made trinucleosomes and mononucleosomes, and recombinant H3 protein. As expected, we detected significant reductions in methylation rates from all substrates in response to RNA binding, with stronger inhibition by the higher-affinity 2G4 RNA (Fig. 3H). The extent of inhibition was limited by the RNA concentration because complete inhibition was achieved with excess 1G4 RNA (Fig. S13A). We also attempted to test the EZH2 R566Y K568Y Q575Y mutant in response to RNA-mediated inhibition (Fig. S13B). However, this mutant had a basal level of activity that was too weak to be accurately quantified, presumably because those mutated residues are responsible for H3 tail loading of PRC2.

### An arginine-rich site of EZH2 binds 1G4 RNA and facilitates RNA-to-DNA hand-off

To examine the candidate sites for 1G4 RNA binding, we generated a subtracted map to focus on the vicinity of the RNA, which was sufficient to visualize one site (EZH2 353-362: KRPGGRRRGR) physically contacting 1G4 (Fig. 4A). This arginine-rich site is part of a disordered loop within the EZH2 SANT2 domain (Fig. 4B), and it has been implicated in binding lncRNA (*24, 35*). Interestingly, a recent study of transcription factors also identified a similar arginine-rich sequence responsible for binding many RNAs (*37*). However, when we tested this interaction by mutagenesis using a truncated EZH2 (EZH2 Δ353-362) and a local charge-reversed EZH2 (EZH2 CR, 353-362: DEPGGEEEGE), we did not detect an obvious reduction of G4 RNA binding (Fig. S14). We then generated an EZH2 double-truncation mutant (EZH2 Δ353-362 Δ494-502) to remove a potentially redundant lysine-rich sequence known to be important for G4-RNA binding (*38*) (Fig. 4B). A 1.5-to-2-fold reduction of binding affinity was detected for this double mutant (Fig. S14E).

**Fig. 4.**
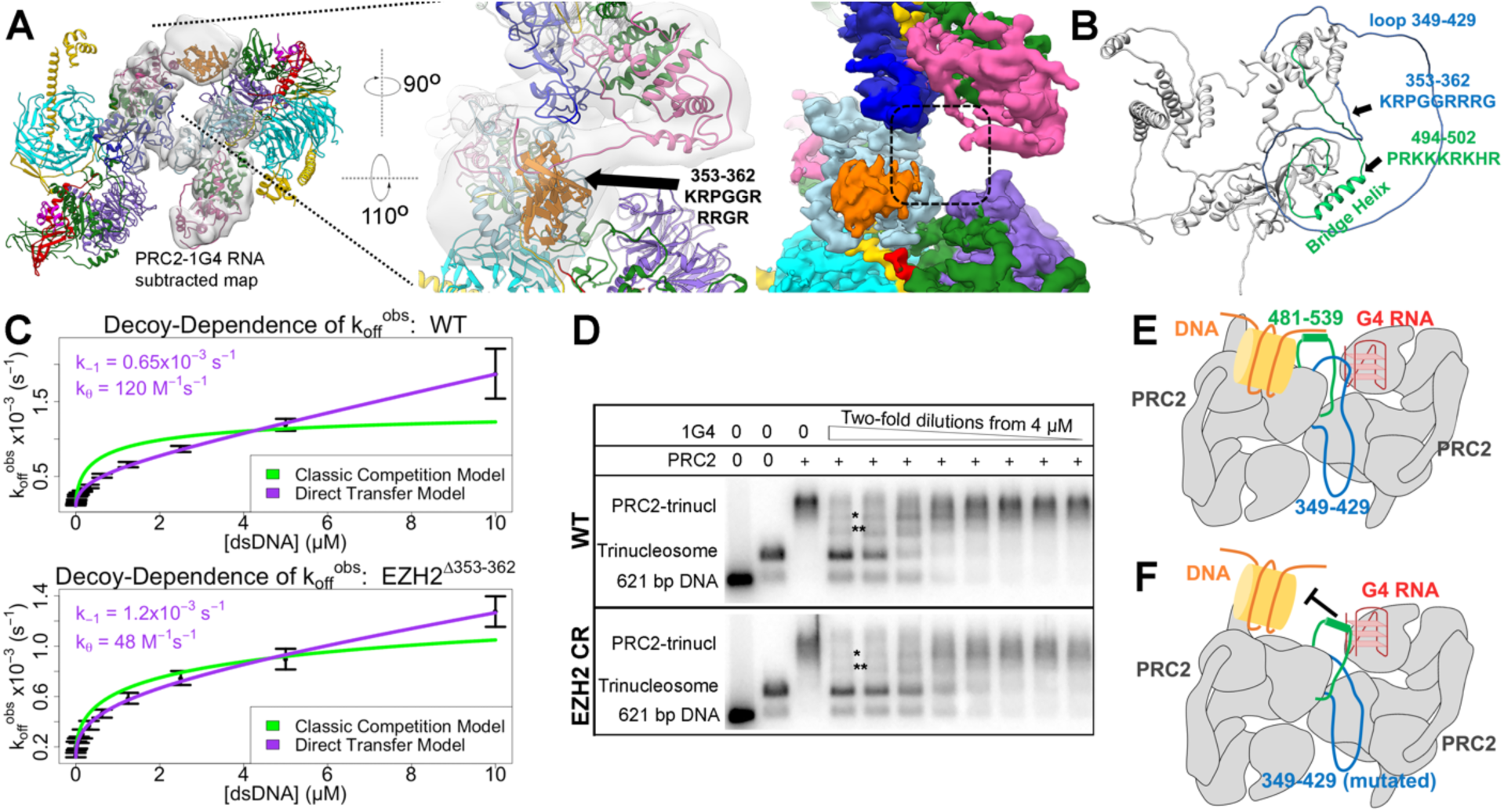
EZH2 353-362 physically contacts G4 RNA and contributes the handoff from RNA to DNA. (**A**) Subtracted map of PRC2-1G4 RNA with zoom-in to emphasize the observed physical interaction of RNA and EZH2 353-362 which was not visualized in high-resolution maps (right). (**B**) EZH2 structure from AlphaFold (*51*) predicts two disordered loops of EZH2. The EZH2 353-362 and EZH2 494-502 are highlighted in blue and green respectively. (**C**) FP assays to monitor the transfer kinetics of PRC2 from fluorescently labeled 1G4 RNA to a dsDNA competitor. FP data were best fit by the model of direct transfer. The ratio of the direct transfer rate constant and the dissociation rate constant (k_θ_/k_-1_) was used to define the propensity of an RNA molecule to direct transfer to DNA (without a free PRC2 intermediate state) relative to classical transfer (releasing from RNA followed by rebinding to DNA). WT PRC2 had k_θ_/k_-1_ = 1.9×10^5^ M^-1^ and EZH2 Δ353-362 mutant had k_θ_/k_-1_ = 0.4 x10^5^ M^-1^. (**D**) Nucleosome-RNA competition assay with radioactive trinucleosome and serial dilutions of 1G4 RNA. (* and **) Incomplete PRC2-trinucleosome complexes as in Fig. 3D. IC_50_ = 1720 ± 560 nM for WT and 773 ± 360 nM for EZH2 CR mutant (Mean ± range of values, N=2). (**E**) Schematic representation of the proposed co-existing intermediate complex of RNA and DNA bound to PRC2. (**F**) Proposed model for EZH2 Δ353-362 and EZH2 CR mutants.

Do residues 353-362 of EZH2 serve a second function in addition to a redundant contribution to RNA binding, especially considering that their sequence is conserved across many species (Fig. S15A)? *Hemphill et al.* recently reported that PRC2 has the intrinsic ability to directly transfer or “hand off” from RNA to DNA, without there ever being a free-enzyme intermediate (*18, 39*).

Such activity could allow RNA to recruit PRC2 to chromatin in cases where the local RNA density is too low to maintain inhibition (*17*). We found that the EZH2 Δ353-362 mutant was five-fold less efficient at direct transfer from RNA to dsDNA (Fig. 4C), indicating that the loop amino acids facilitate direct transfer. Mutating the 353-362 loop also increased RNA competition with trinucleosomes in an equilibrium binding experiment (Fig. 4D) and increased RNA inhibition of PRC2 HMTase activity (Fig. S15B). In all cases, mutating the 353-362 loop favored RNA over DNA/nucleosome linker binding.

Therefore, we propose a model (Fig. 4E and F) in which EZH2 retains an arginine-rich RNA-binding 353-362 site and a lysine-rich DNA-binding 494-502 site to enable a transient intermediate with both RNA and DNA bound, which allows it to hand off from RNA to DNA or to nucleosome linkers. Due to the sequence and spatial proximity of those two sites, 494-502 can likely compensate for RNA binding in the EZH2 Δ353-362 and EZH2 CR mutants.

## Discussion

In the past ten years, lncRNAs and pre-mRNAs have become prominent in discussions of PRC2 regulation (*4, 5*). Because of the flexible nature of PRC2-RNA interactions (*12*), deciphering molecular details has been challenging and has lagged behind functional observations. Here, we utilized a PRC2-preferred G4 RNA, which is representative of the broad PRC2 transcriptome (*15*), and a biotinylated RNA-streptavidin grid method to determine the cryo-EM structure of an RNA-bound PRC2 complex. This structure agrees with the earlier conclusion that DNA and RNA binding are mutually antagonistic (*20, 23*), but it provides a much more interesting mechanism than just competition on overlapping sites. Instead, G4 RNA triggers formation of a PRC2 dimer that occludes the DNA-binding amino acids. Based on the present structure and biochemical and biophysical data, we propose a model that can reconcile seemingly contradictory ideas (Fig. 5), as follows.

**Fig. 5.**
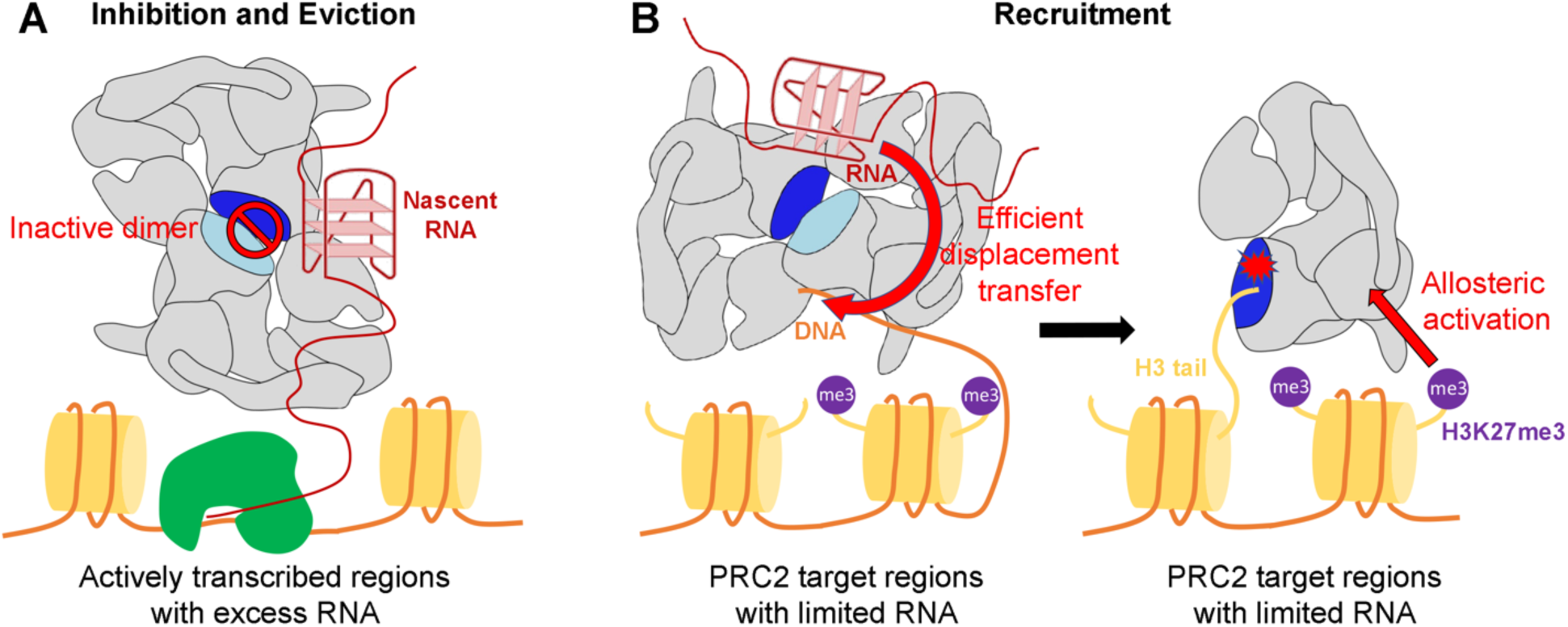
Suggested roles of G4-forming RNA in PRC2 regulation. (**A**) PRC2 is evicted from active loci as an inhibited dimer. (**B**) After recruitment to its target loci by RNA, PRC2 engages in a transient complex with both RNA and nucleosome DNA to facilitate transferring onto the chromatin, accompanied by dissociation of the dimer. Finally, H3K27me3 binds EED to activate PRC2 for histone methylation.

First, actively transcribed loci, which need to avoid silencing by PRC2, generate nascent RNA transcripts that dimerize PRC2. The dimer simultaneously inactivates two PRC2 complexes (no H3 tail binding) and evicts them from local chromatin (no nucleosome DNA binding) (*14, 21, 22*), which prevents a single non-specific H3K27me3 or JARID2-116me3 from allosterically activating PRC2 and productively amplifying the methylation across nearby regions (*30, 40, 41*). Our structure shows that residues of the EZH2 CXC domain, known to load the H3 tail into the catalytic groove, are occupied in a protein-protein interaction to reinforce the dimerization. Second, although PRC2 recruitment remains controversial (*42*), PRC2 has the intrinsic ability to translocate onto target chromatin as it simultaneously dissociates from inhibitory RNA. A recent study genetically defined the importance of RNA interaction to promote PRC2 target chromatin occupancy (*17*). Consistent with this, we find our G4-mediated PRC2 dimer is reversible. EZH2 maintains two conserved but redundant sites for forming a transient intermediate in which two nucleic acids are simultaneously bound; this could facilitate efficient direct transfer from RNA to DNA (*18, 39*), and therefore, enhance chromatin occupancy. Finally, after recruitment, existing H3K27me3 can interact with EED to serve as an allosteric activator for PRC2 activity (*30, 43*).

Our structure describes one mode of RNA recognition by PRC2, but there may very well be others. Other reported RNA-binding sites including the RNA-binding region (RBR) adjunct to the bridge helix of EZH2 (residues 494-502) (*38*), the stimulatory recognition motif (SRM) of EZH2 (residues 127-153) (*24*), the EED amino acids close to EZH2 SRM (residues 336–355) (*24*), and the JARID2 RBR (residues 332–358) (*44*). Although we did not obtain any subclass maps having a clear RNA density in proximity to those regions, this is not sufficient to reject their RNA binding potential, especially because we did not completely abolish RNA binding in our mutagenesis studies.

PRC2 has been shown to dimerize without RNA binding. A four-subunit PRC2 holoenzyme was reported as a dimer in solution at high complex concentration (*45*). Because the structure of this dimer remains unknown, we do not know if it is related to the RNA-mediated dimer described here. A small fraction of six-subunit PRC2.2 intrinsically dimerized with an instinct dimer arrangement (Fig. S16). In addition, two reported domain-swapped PRC2 dimers -- PRC2-PCL and PRC2:EZH1(*28, 31*)--have completely different architectures than our RNA-mediated dimer.

Besides PRC2, many chromatin-associated complexes have been found to interact with RNA molecules, including other histone modifiers (*2*), transcription factors (*37, 46*), RNA polymerase (*47, 48*), and DNA methyltransferase (*49, 50*). Like PRC2, they tend not to have canonical RNA-recognition motifs and bind RNA broadly, and obtaining molecular structures of the RNA-protein complexes has been very challenging. Ultimately, solving additional structures of RNA bound to epigenetic modifiers will reveal mechanisms by which RNA serves direct regulatory roles, rather than simply serving as a messenger.

## Acknowledgments

We thank R. Yan, Z. Yu, and S. Yang (Janelia Cryo-EM Facility), C. Moe (University of Colorado Boulder BioKEM Facility), and G. Morgan (University of Colorado Boulder Electron Microscopy Service) for microscope operation and data collection. We thank L. Yao for help with streptavidin-affinity grid preparation; A. Iragavarapu for help with plasmid preparation; R. Fenske for help with FP assay; A. Erbse (University of Colorado Boulder Shared Instruments Pool) for help with MST assay. We thank Y. Long (Weill Cornell Medicine) and C. Lim (University of Wisconsin Madison) for helpful discussions. We thank B. Greber for helpful suggestions related to model refinement.

## Author contributions

Conceptualization: J.S., V.K., and T.R.C. Investigation: J.S., A.R.G., W.O.H., V.K., and T.R.C. Visualization: J.S., A.R.G., W.O.H., and V.K.

## Funding acquisition

V.K. and T.R.C. Project administration: V.K. and T.R.C. Supervision: V.K. and T.R.C. Writing – original draft: J.S., V.K., and T.R.C. Writing – review & editing: all authors.

## Data and materials availability

All data needed to evaluate the conclusions in this paper are present either in the main text or in the supplementary materials. Cryo-EM density maps and fitted models have been deposited in the Electron Microscopy Data Bank (EMD-29578, consensus map) (EMD-29647, Body1 from multibody refinement) (EMD-29656, Body2 from multibody refinement) and the Protein Data Bank (PDB: 8FYH). Requests for resources, reagents, plasmids, and cell lines used in this study and additional information should be directed to corresponding authors V.K. and T.R.C.

## Supplementary Materials

### Materials and Methods

#### Protein expression and purification

Full-length EED, SUZ12, RBAP48, His-tagged EZH2 isoform 2, Strep-GFP-tagged embryonic isoform of AEBP2, and Strep-GFP-tagged truncated JARID2 (amino acids 119-450) were assembled into a single multi-bac plasmid. This multi-bac plasmid was used to make infectious baculovirus stock in Sf9 cells using the Bac-to-Bac system (Invitrogen). To express recombinant complex, HighFive cells were transfected with baculovirus at 28°C for 66 hours and frozen in liquid nitrogen until use.

All purification steps were performed in a 4°C cold room. Cells were lysed in lysis buffer (25 mM HEPES pH 7.9 at 4°C, 250 mM NaCl, 2 mM MgCl_2_, 1 mM TCEP, 10 mM imidazole, 0.5% NP-40, 10% glycerol, and protease inhibitor cocktail) for 1 hour and sonicated with mild strength. Debris was then removed by centrifugation at 15,000 rpm for 35 minutes. The supernatant was incubated with Ni-NTA agarose resin (Qiagen) for 1 hour, and resin was washed with 10 column volumes (CV) of lysis buffer, 10 CV of high-salt wash buffer (25 mM HEPES pH 7.9 at 4°C, 1 M NaCl, 2 mM MgCl_2_, 1 mM TCEP, 0.01% NP-40, and 10% glycerol), and 20 CV of low-salt wash buffer (25 mM HEPES pH 7.9 at 4°C, 150 mM NaCl, 2 mM MgCl_2_, 1 mM TCEP, 30 mM imidazole, and 10% glycerol). Proteins were then eluted in elution buffer (25 mM HEPES pH 7.9 at 4°C, 150 mM NaCl, 2 mM MgCl_2_, 1 mM TCEP, 300 mM imidazole, and 10% glycerol) and dialyzed for 2 hours in buffer (25 mM HEPES pH 7.9 at 4°C, 150 mM NaCl, 2 mM MgCl_2_, 1 mM TCEP, and 10% glycerol) to remove imidazole. After concentrating to 3-5 mg/ml, proteins were incubated with TEV protease overnight. We used an AKTA-FPLC system for subsequent purification with a HiTrap Heparin HP column (Cytiva) and a Superose 6 increase 10/300 column (GE Healthcare). Heparin column was equilibrated with buffer I (20mM HEPES pH 7.9 at 4°C, 150 mM NaCl, 2 mM MgCl_2_, 1 mM TCEP, and 10% glycerol), and sample was eluted with a linear gradient of buffer II (20mM HEPES pH 7.9 at 4°C, 2 M NaCl, 2 mM MgCl_2_, 1 mM TCEP, and 10% glycerol). The Superose 6 increase column was equilibrated and performed with final storage buffer (25 mM HEPES pH 7.9 at 4°C, 150 mM KCl, 2 mM MgCl_2_, 10% glycerol, and 1mM TCEP). Protein complex was flash frozen in liquid nitrogen as single-use aliquots and stored at −80°C.

For preparations of mutated PRC2 complexes, Q5 site-directed mutagenesis (NEB) and plasmid synthesis (GenScript) were used to generate EZH2 constructs with corresponding mutations. Then NEBuilder HiFi DNA assembly (NEB) was applied to assemble the final multi-bac plasmids with mutated EZH2.

#### RNP complex assembly

We purchased two G-quadruplex (G4)-forming RNA oligos, 1G4 and 2G4, from IDT RNA oligonucleotide synthesis including HPLC purification service. 1G4 is a 50-nt single-G4 RNA with a 5’ biotin modification, followed by a 30-nt A-rich linker sequence to provide flexibility between biotin and the functional G4 group. 1G4 has (GGGAA)_4_ sequence which folds into a stable G4 structure. 2G4 has two-independent-G4 motifs separated by a 15-nt A-rich linker, 85 nt total. A-rich sequences were chosen for the linker because poly(A) RNA does not interact with PRC2 (*15*). G4 RNA was heated at 95°C for 3 min, snap-cooled on ice for 5 min, refolded in RNP complex buffer (25 mM HEPES pH 7.9 at 4°C, 50 mM KCl, 2 mM MgCl_2_, 10% glycerol, and 1mM TCEP) at 37°C for 30 min. PRC2 and S-adenosyl homocysteine (SAH) were added into the reaction at final concentration of 600 nM and 40 µM correspondingly, and the reaction was incubated at 30°C for 30 min to assemble the RNP complex.

#### Cryo-EM sample preparation

Quantifoil Au 2/2 streptavidin-affinity grids were made in-house using procedures previously described (*32*). Grids were re-hydrated by EM preparation buffer I (25 mM HEPES pH 7.9 at 4°C, 50 mM KCl, 2.5% glycerol, and 1mM TCEP) at room temperature (RT) for 1 hour. After removing remaining buffer, 4 µl of the assembled RNP complex was applied onto streptavidin-affinity grids and incubated for 5-10 min in a humidified chamber. The grid was washed with 40 µl of EM preparation buffer I and EM preparation buffer II (25 mM HEPES pH 7.9 at 4°C, 50 mM KCl, 2.5% glycerol, 0.01%NP-40, and 1mM TCEP). After the washes, the buffer was wicked away using Whatman filter paper, and 4 µl of the EM preparation buffer II was added immediately. The grid was then transferred to the Leica EM GP2 plunge freezer and blotted for 2-3 s at 8°C and 90% humidity, and then plunged into liquid ethane.

Negative staining of streptavidin-affinity grids was applied through the same protocol. Instead of using a plunge freezer, 5 droplets of 40 µl stain was used. For negative staining using continuous carbon grids, 4 µl of 100 nM protein was applied on glow-discharged carbon film 400 mesh Cu grid and incubated for 20 s before staining.

#### Cryo-EM data collection and processing

Cryo-EM dataset was collected on a Titan Krios equipped with Gatan K3 direct detector in super-resolution mode and a Cs-corrector. A GIF quantum energy filter was used for collection with a 20-eV slit width. Movies were recorded at a nominal magnification of 81,000x, corresponding to a calibrated pixel size of 0.844Å (super resolution 0.422 Å). Data acquisition was performed using SerialEM for automated data collection with a defocus range of −2.0 to −0.6 µm. The total dose for our dataset was 60 electrons per square angstrom (e^−^/Å^2^). It was acquired as dark-subtracted, non–gain corrected movies, and gain correction was applied during motion correction using MotionCor2 (*52*).

Negative stain samples were collected on a Tecnai F20 microscope operated at 200 kV, with a Gatan K3 direct detector, at a nominal magnification of 25,000x corresponding to 1.449 Å per pixel, using a dose of 20-50 e^−^/Å^2^. Data was processed in RELION 4 (*53*). The movie frames were aligned using MotionCor2 (*52*) and CTF parameters were fit using CTFFIND (*54*). The background streptavidin lattice of each micrograph was subtracted using in-house scripts (*32*). LoG automatic picking was applied to pick individual particles. Initial models were generated within RELION from negative staining data and used as reference for the first round of 3D classification. Subsequent processing steps including several runs of regular 3D classification and 3D classification without alignment (regularization parameter T=24), which used references from previous good classes. The selected 217,196 particles were then re-extracted and subjected to per-particle defocus refinement, beam-tilt refinement, 3D refinement, and postprocessing to generate the consensus map. Soft-edged masks used in multibody refinement (*33, 55*) were generated within RELION. Local resolution estimation was performed using the same soft, spherical masks used during refinement.

#### Model building

Individual PRC2 protomers were built using cryo-EM maps from the multibody refitment. The coordinates of nucleosome-bound PRC2 six-subunit complex (PDB: 6WKR) (*26*) was used as a starting model from which all the coordinates were adjusted and rebuilt in the new map using COOT (*56*). We used EZH2 isoform 2 (297-298:HP→HRKCNYS) in this study, which has a five-residue insertion that forms an unstructured loop, instead of the more frequently used isoform 1. This insertion only extended the length of its belonging loop without altering other defined secondary and tertiary structures of EZH2 (Fig. S3C). To allow comparison with published PRC2 structures, EZH2 residue numbers in this study are corresponding to isoform 2. Region corresponding to EZH2 isoform 2 was built de novo into the EM density in COOT. 1G4 model was adapted from PDB: 2M18, and then docked into our map for the position of clear RNA density. We ignored the very weak RNA density. The model of individual PRC2 promoter was subjected to global refinement and minimization in real space using PHENIX (*57*). These were then subjected to manual inspection and adjustment in COOT followed by refinement again in PHENIX. The models of the individual PRC2 were then refined in PHENIX against the full consensus map with local grid search to validate the interface rotamers (*58*). The cryo-EM density maps and the molecular graphics were prepared with Chimera and ChimeraX (*59*). The distance between PRC2 protomers was calculated using residues from EZH2 SANT1 domain and SUZ12 RRM-like domain.

#### Analytic size-exclusion chromatography

In a 50 µl reaction, PRC2 and refolded G4 RNA were mixed at a final concentration of 2 µM with RNP complex buffer (25 mM HEPES pH 7.9 at 4°C, 50 mM KCl, 2 mM MgCl_2_, 10% glycerol, and 1mM TCEP). The reaction was incubated at 30°C for 30 min to complete RNP assembly, and then injected into a Superose 6 increase 3.2/300 column (Cytiva) pre-equilibrated with the RNP complex buffer. The column was run at a flow rate of 0.02 ml/min, monitored by UV260 and UV280 detectors. Gel filtration standard (Bio-rad) was injected and run at the same protocol to estimate the molecular weight of unbound PRC2 and G4-bound RNP complexes.

#### Microscale Thermophoresis (MST)

We purchased 3’ Cys5-labeled RNA oligos, 1G4 and (GA)_20_, from IDT RNA oligonucleotide synthesis including HPLC purification service. RNA oligos were heated, snap-cooled, and refolded as indicated in RNP assembly. Our MST instrument is a Nano-BLUE/RED Monolith NT.115 from NanoTemper Technologies, equipped with two LED-filter combinations. We followed manufacturer’s instructions for our experiments. Briefly, 20 nM RNA with serial dilutions of PRC2 proteins were incubated in MST assay buffer (10 mM HEPES pH 7.9 at 4°C, 50 mM KCl, 2 mM MgCl_2_, 0.5 mg/ml BSA, and 1mM TCEP) at 30°C for 15 min prior to measurement. Lithium chloride MST assay buffer (10 mM HEPES pH 7.9 at 4°C, 50 mM LiCl, 2 mM MgCl_2_, 0.5 mg/ml BSA, and 1mM TCEP) was substituted in particular reactions. The graphic plots were generated from the default evaluation software provided by the equipment.

#### Electrophoretic mobility shift assay (EMSA)

EMSA was conducted as previously described (*38*) with modifications. 1G4 and 2G4 RNA oligos were purchased from IDT RNA oligonucleotide synthesis without 5’ biotin modification. After 5’ end-labeling with gamma-^32^P-ATP, oligos were heated, snap-cooled, and refolded in EMSA binding buffer (50 mM Tris-HCl pH 7.5 at 25°C, 100 mM KCl, 2.5 mM MgCl_2_, 0.1 mM ZnCl_2_, 2 mM 2-mercaptoethanol, 0.1 mg/ml BSA, 0.1 mg/ml fragmented yeast tRNA, and 5% glycerol). Protein samples were diluted and incubated with refolded RNA at 30°C for 30 min. Sample was loaded to 1% agarose gel (SeaKem GTG Agarose) buffered with 1X TBE and resolved at 66 V for 90 min in a 4°C cold room. Gels were vacuum dried for 60 min at 80°C. Dried gels were exposed to phosphorimaging plates and signal acquisition was performed using a Typhoon Trio phosphorimager (GE Healthcare). Signal intensities were quantified by ImageQuant TL. Data were plotted to calculate Kd values using the Prism software.

#### Fluorescence polarization (FP)

Regular FP assays (binding affinity measurement) and modified FP assays (competitive dissociation measurement) were performed as previously described (*18*). Briefly, in the FP binding assays, 5 nM RNA or DNA was incubated with serial dilutions of six-subunit PRC2 in FP buffer (50 mM TRIS pH 7.5 at 25°C, 25 mM KCl, 2.5 mM MgCl_2_, 0.1 mM ZnCl_2_, 0.1 mg/mL BSA, 5% glycerol, and 2 mM 2-mercaptoethanol) for 30 min at RT. Fluorescence polarization readings were then taken for 30 min in 30 s intervals with a TECAN Spark microplate reader. Raw data were analyzed in R v4.1.1 with FPalyze (github.com/whemphil/FPalyze). Hill model included the Hill coefficient in the fitted equation to describe cooperativity. In the modified FP assays, 5 nM RNA and 100 nM PRC2 were incubated in a pre-reaction mix to achieve binding equilibrium. Competitive dissociation reactions were initiated by addition of varying concentrations of DNA decoy to the corresponding pre-reaction mix, then fluorescence polarization readings were immediately taken at 25°C for 120 min in 30 s intervals with a TECAN Spark microplate reader. Raw data were analyzed in R v4.1.1 with FPalyze.

#### Methyltransferase activity assay

For reaction time-based assays, 400 nM PRC2 and 400 nM refolded G4 RNAs were pre-incubated at 30°C for 30 min to reach binding equilibrium, and then assembled into methyltransferase reaction mix including 1X methyltransferase buffer (25 mM HEPES pH 7.9 at 4°C, 50 mM KCl, 2 mM MgCl_2_, 10% glycerol, and 1mM TCEP), 0.1 mg/ml BSA, 1X protease inhibitor, 1 µl/20 µl RNase inhibitor, 10 µM ^14^C SAM (PerkinElmer), and 0.16 mg/ml polynucleosome or 200 nM trinucleosome or 250 nM mononucleosome or 6.5 µM recombinant H3 (NEB). Samples were collected at 0, 3, 6, 10, 15, 20, 30, 45, and 60 min from the same tube incubating at 30°C. Proteins were separated through NuPAGE 4-12% gel (Invitrogen) by running at 180V for 52 min. Gel was vacuum dried at 80°C for 30 min, and then exposed to phosphorimaging plates. Signal intensities were quantified by ImageQuant TL and plotted by Microsoft Excel.

For RNA concentration-based assays, 400 nM PRC2 and serial dilutions of refolded 1G4 RNA were pre-incubated and assembled into methyltransferase reaction mix including 1X methyltransferase buffer, 0.1 mg/ml BSA, 1X protease inhibitor, 1 µl/20 µl RNase inhibitor, 10 µM ^14^C SAM (PerkinElmer), and 200 nM trinucleosome. Reactions were performed at 30°C for 20 min. NuPAGE running, signal acquisition and quantification were same as previous.

#### Nucleosome-RNA competition assay

We prepared ^32^P-labeled trinucleosomes in-house by adding a small fraction of radiolabeled DNA into non-labeled DNA during trinucleosome assembly. 600 nM PRC2, 150 nM labeled trinucleosome, and serial dilutions of refolded 1G4 RNA were combined in RNP complex buffer (25 mM HEPES pH 7.9 at 4°C, 50 mM KCl, 2 mM MgCl_2_, 10% glycerol, and 1mM TCEP) at 30°C for 30 min. Sample was loaded to 1% agarose gel (SeaKem GTG Agarose) buffered with 1X TBE and resolved at 66 V for 110 min in a 4°C cold room. Gels were vacuum dried for 60 min at 80°C. Dried gels were exposed to phosphorimaging plates and signal acquisition was performed using a Typhoon Trio phosphorimager (GE Healthcare). Signal intensities were quantified by ImageQuant TL. Data were plotted to calculate IC_50_ values using the Prism software. Incomplete nucleosom-PRC2 complexes were included into the bound fraction and free DNA were included into the unbound fraction.

---

**Fig. S1.**
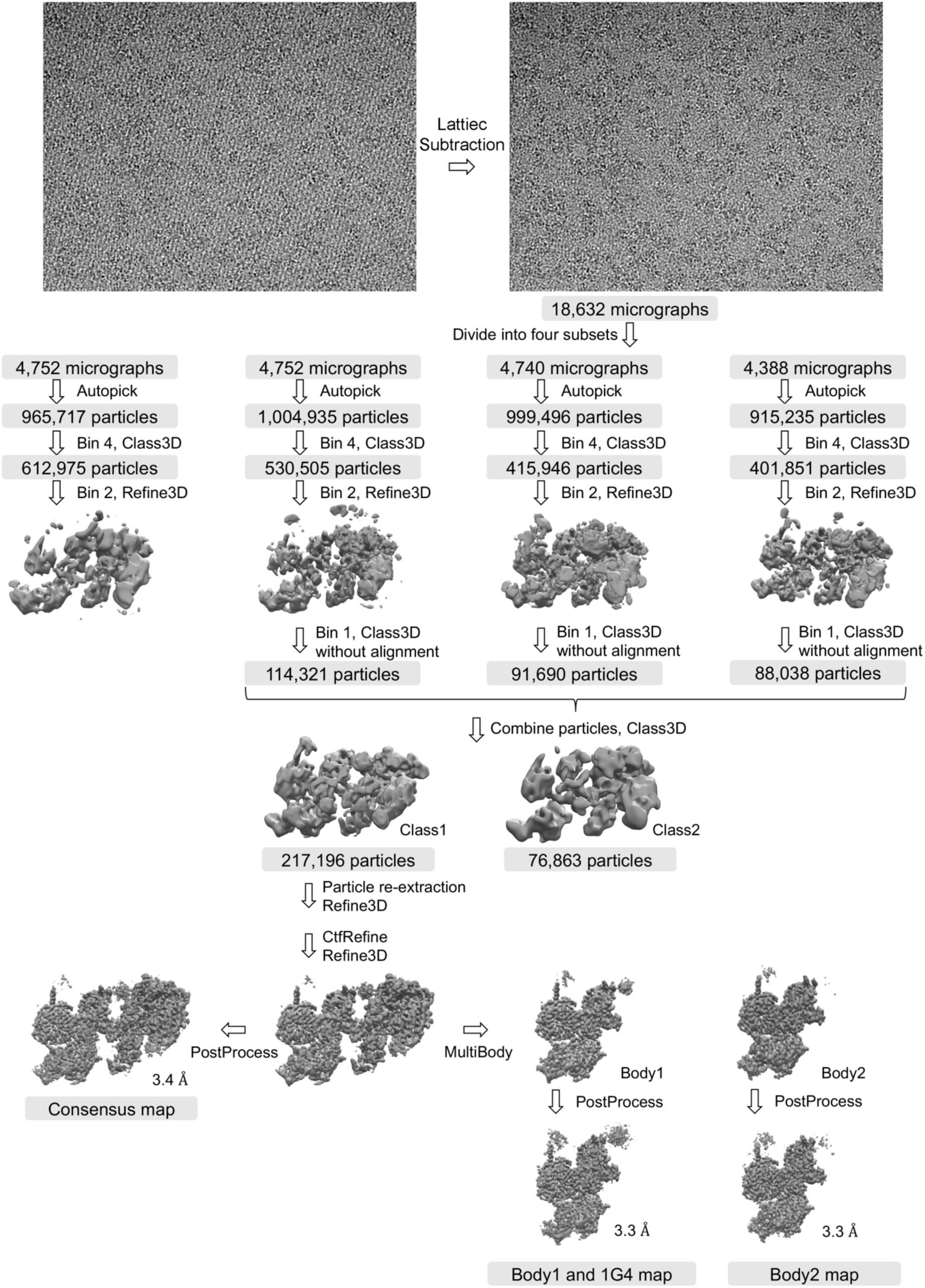
Single-particle cryo-EM image processing workflows for PRC2-1G4 RNA complex.

**Fig. S2.**
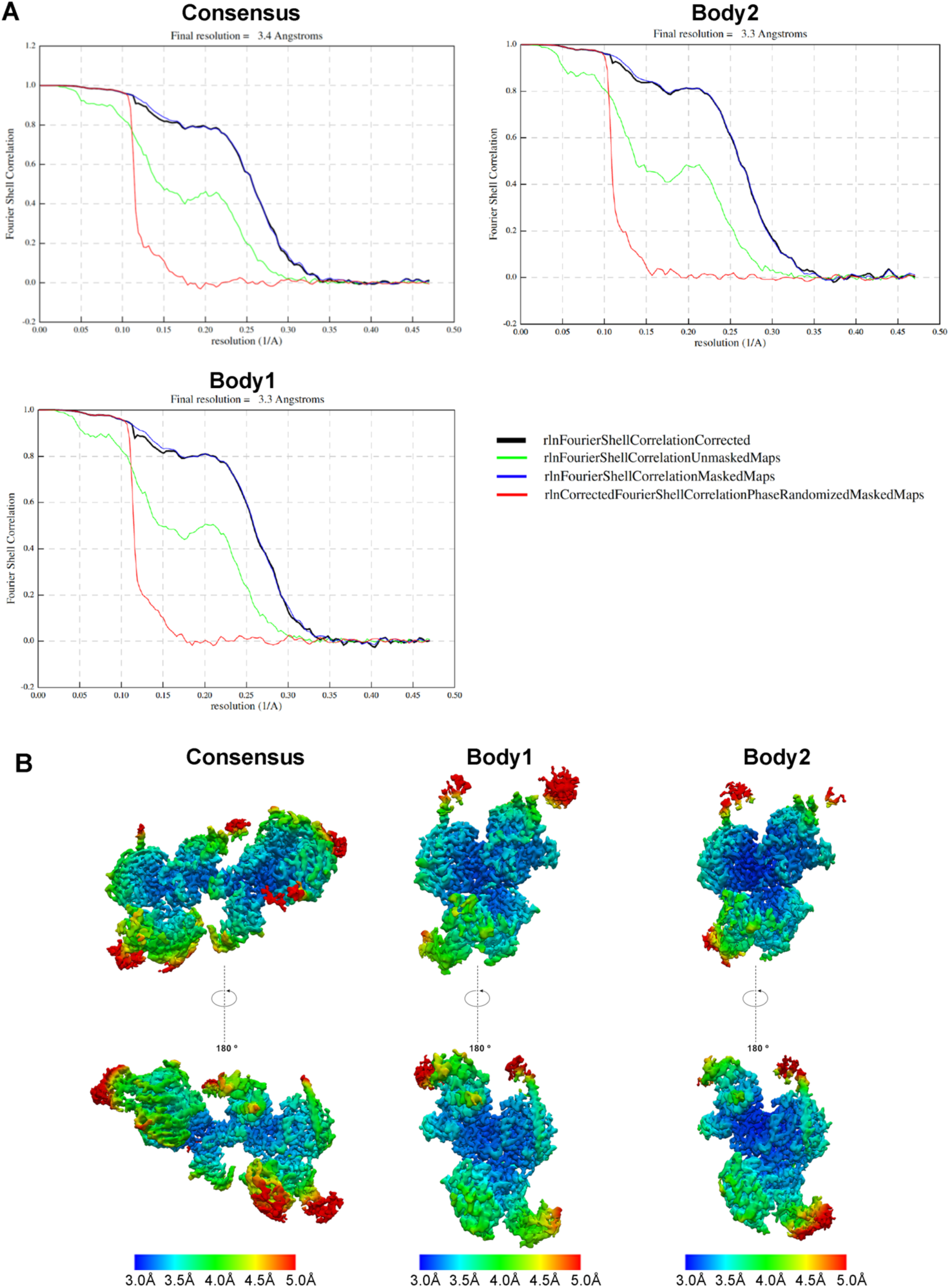
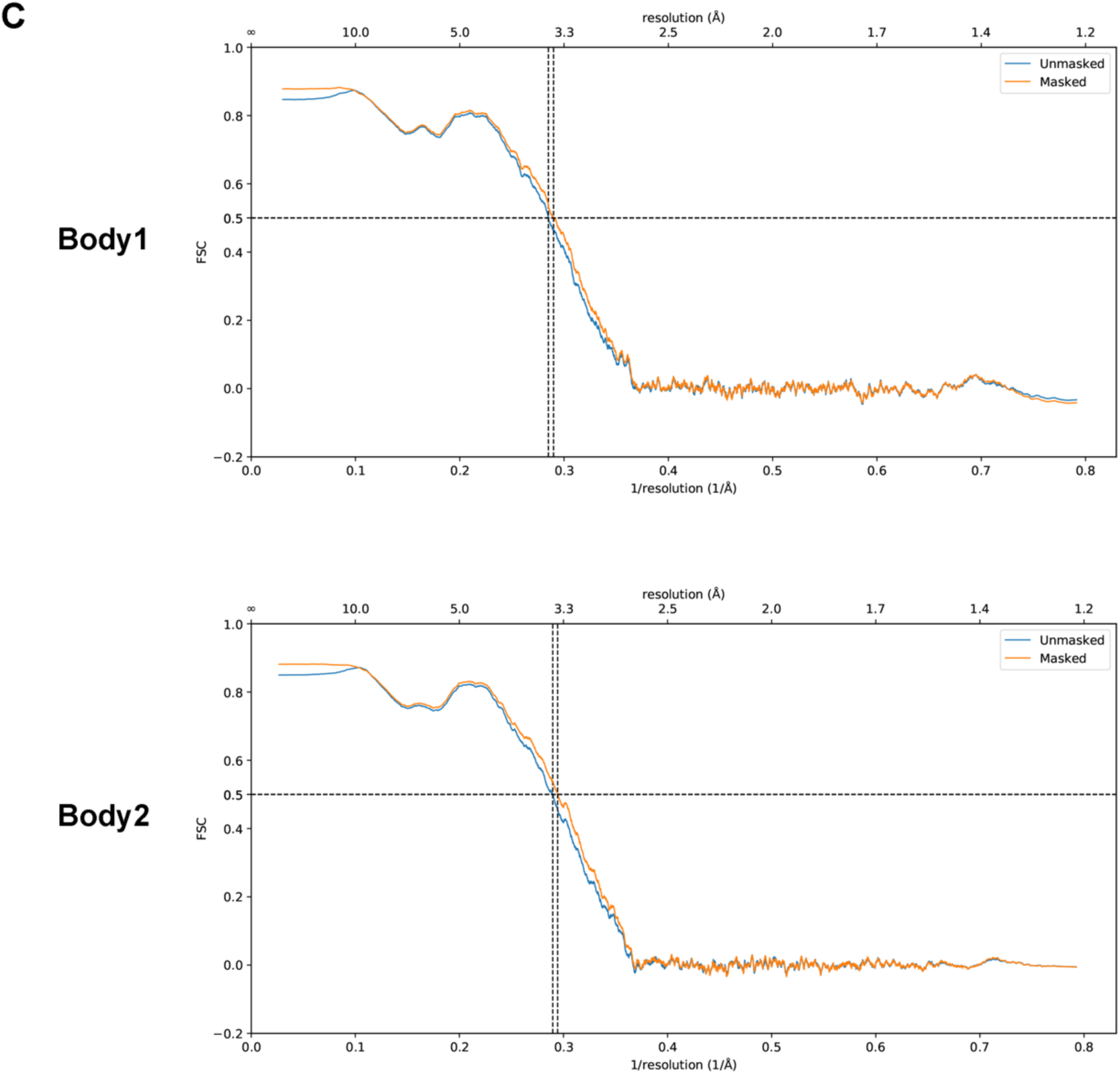
Cryo-EM map analysis. (**A**) Fourier shell correlation curves for consensus map and maps of two individual bodies. (**B**) Local-resolution density maps of consensus and two bodies. (**C**) Model vs Map FSC for Phenix-refined models of two bodies.

**Fig. S3.**
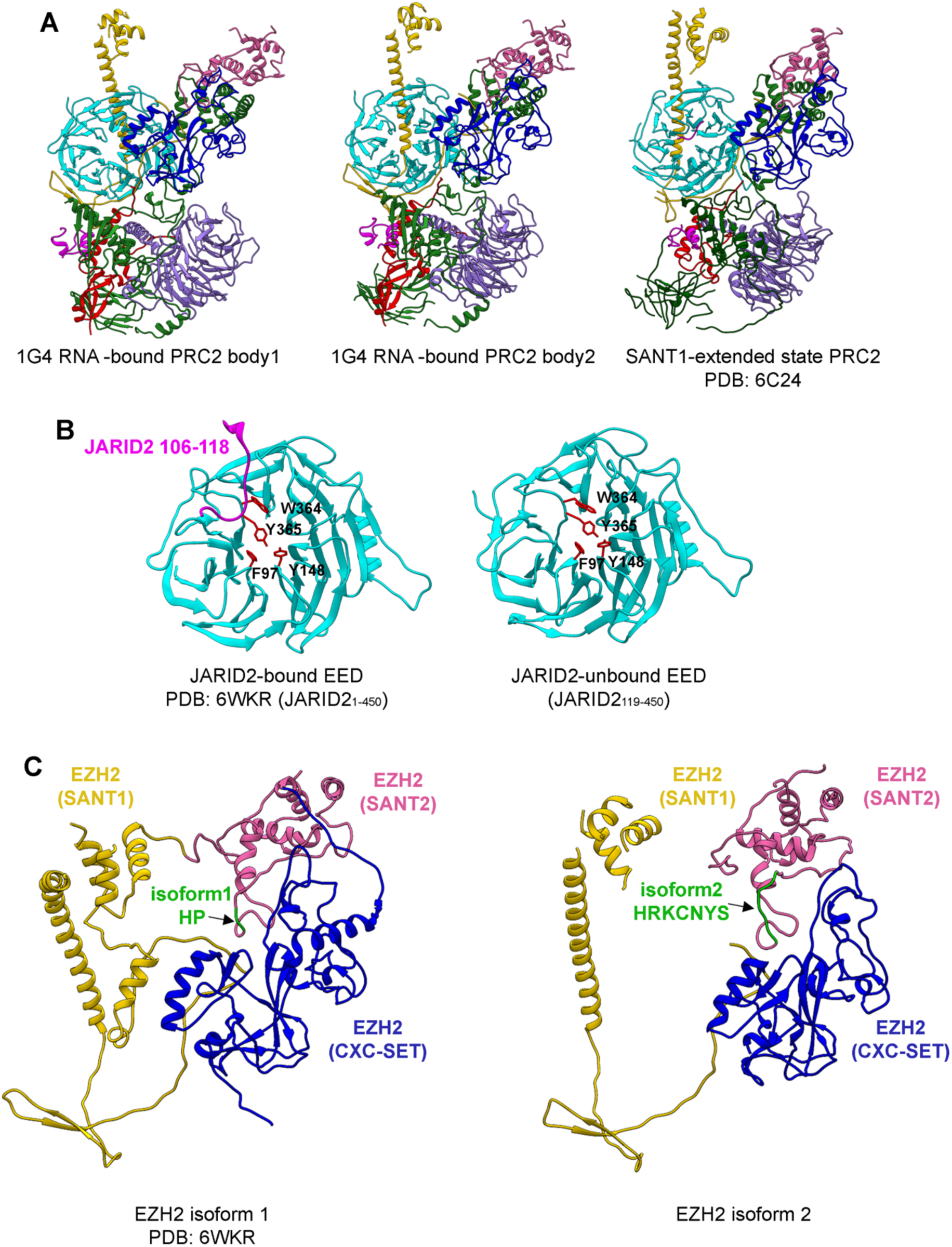
Structure comparisons. (**A**) Overall subunit arrangements of RNA-bound PRC2 body1, body2 and SANT1-extended state PRC2 are nearly identical. EZH2 (SANT1) is highlighted in gold, EZH2 (SANT2) in hot pink, EZH2 (CXC-SET) in blue, EED in cyan, RBAP48 in purple, SUZ12 in green, JARID2 in magenta, and AEBP2 in red. (**B**) Structure comparison of JARID2-bound EED and JARID2-free EED. We truncated JARID2 N-terminus (1–118) to avoid JARID2-RNA competition suggested by others (*24*). The N-terminal region of JARID2 contains an EED binding site (residues 106-118 with K116me3) functioning as a H3K27me3 mimic and PRC2 activator (*40*). Although our JARID2_119-450_ lacks it, EED does not experience any major conformational change. EED and JARID2 are colored in cyan and magenta respectively. EED residues F97, Y148, W364 and Y365 have been proposed for JARID2 association and highlighted in red stick presentation here. (**C**) Structure comparison of EZH2 isoform 1 and isoform 2 (297-298:HP→HRKCNYS) showing the same secondary and tertiary arrangements. EZH2 isoform 1 structure is adapted from nucleosome-bound PRC2 (PDB:6WKR). SANT1 domain is highlighted in gold, SANT2 in hot pink, CXC-SET in blue, and different residues between isoforms in green.

**Fig. S4.**
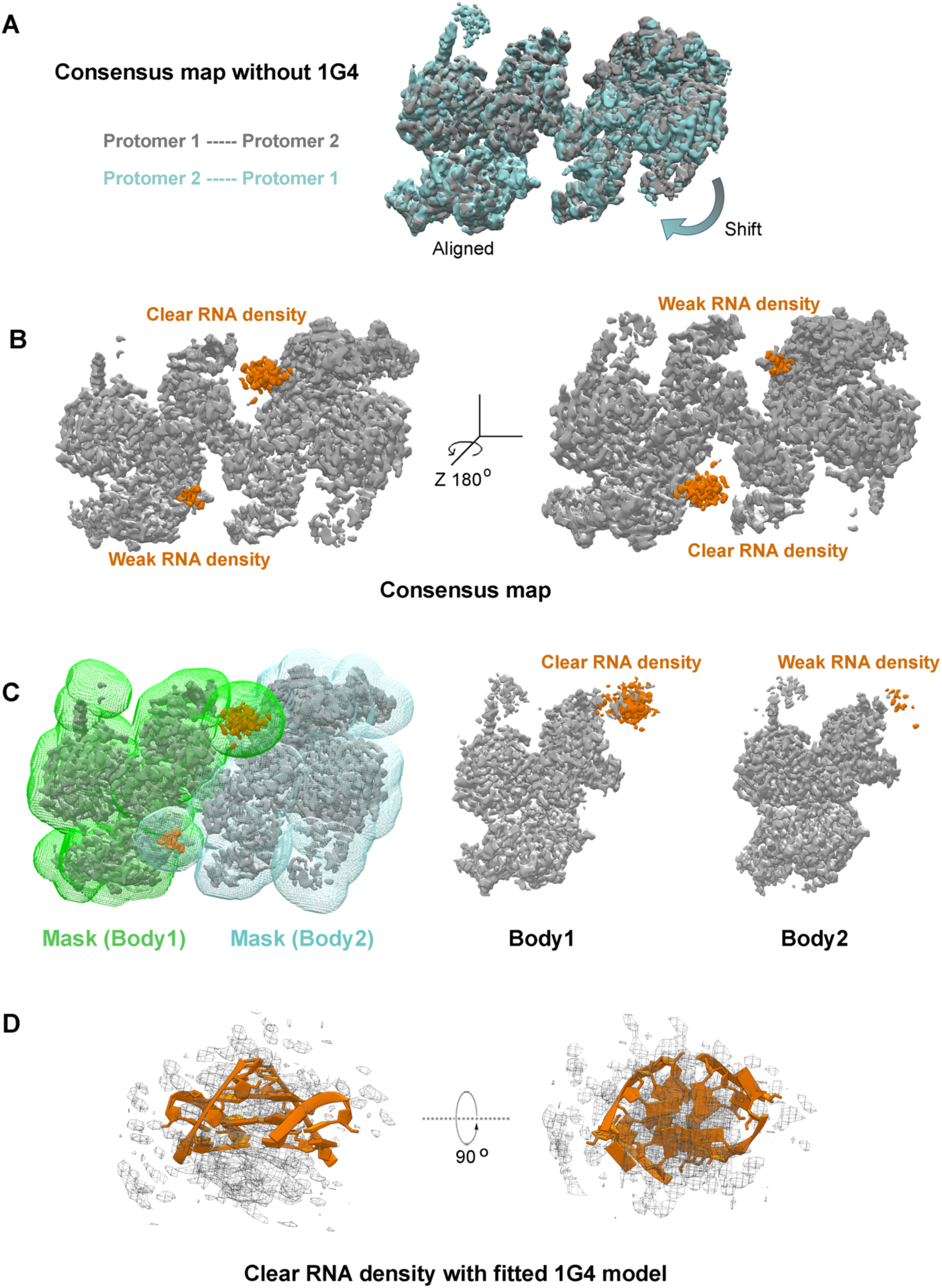
PRC2-1G4 RNP has imperfect C*_2_* symmetry. (**A**) The consensus map without 1G4 density was duplicated and rotated 180 degrees cross the Z-axis. The protomer 1 of original map (gray) is aligned with the protomer 2 of the rotated map (cyan) to highlight the imperfect symmetry. (**B**) A clear RNA density and a weak density on two symmetric sites were distinguished from the PRC2-1G4 consensus map. (**C**) We grouped one PRC2 promoter with clear RNA density into Body1 and the other promoter with weak density into Body2 in the multibody refinement. Two masks are highlighted in green and blue, respectively. After refinement, the weak density was discontinuous. (**D**) 1G4 model (adapted from PDB:2M18) was fitted into the clear RNA density (mesh).

**Fig. S5.**
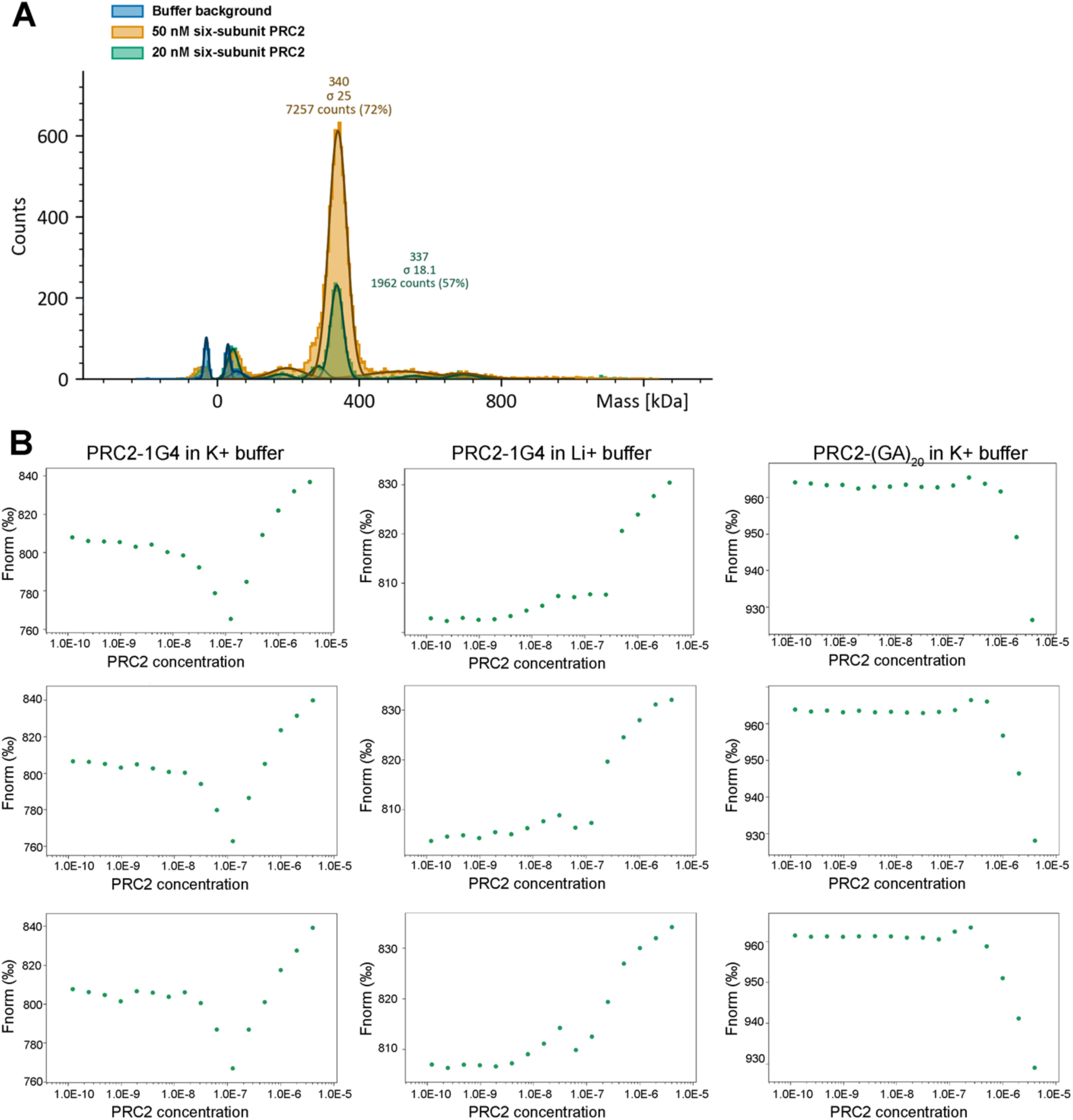
PRC2 six-subunit complex is a monomer in solution and dimerizes by G4 RNA binding. (**A**) Mass photometer results of six-subunit PRC2 at two protein concentrations. (**B**) MST profiles of PRC2-RNA interactions. The fluorescence change in MST signal is normalized (Fnorm), defined as F_hot_/F_cold_ (F_hot_ as the hot region at 20 s after IR laser heating and F_cold_ as the cold region at 0 s) (*60*). Reaction was carried in the same K+ buffer as used in the cryo-EM. Li+ destabilizes G4 structure (*61*) to serve a negative control (middle). (GA)_20_ weakly interacts with PRC2 and also serves a negative control (right).

**Fig. S6.**
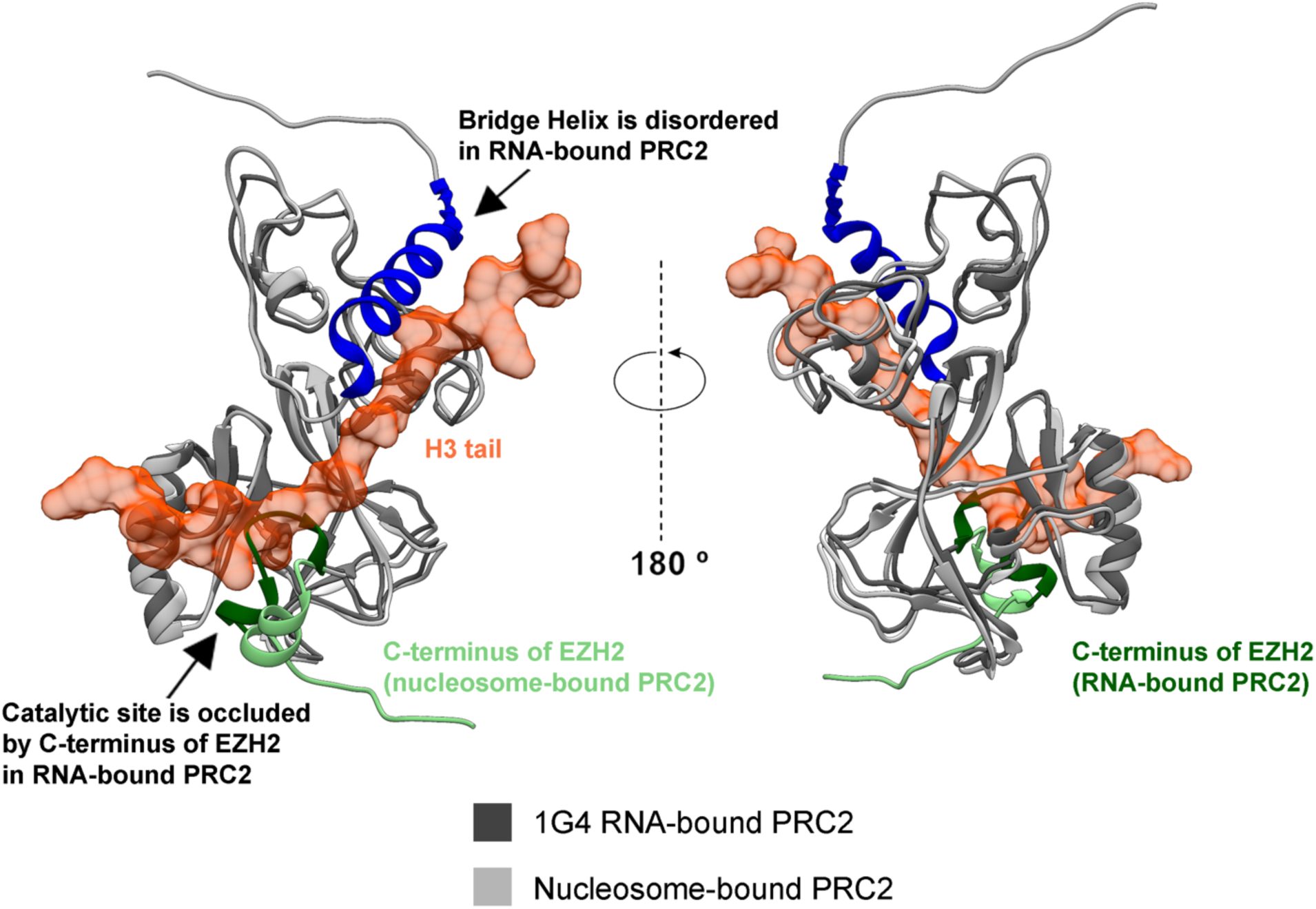
Superimposed comparison of EZH2 CXC and SET domains of nucleosome-bound and RNA-bound PRC2. Two differences are distinguished and highlighted with arrows. Nucleosome H3 tail is shown in orange space-filling model.

**Fig. S7.**
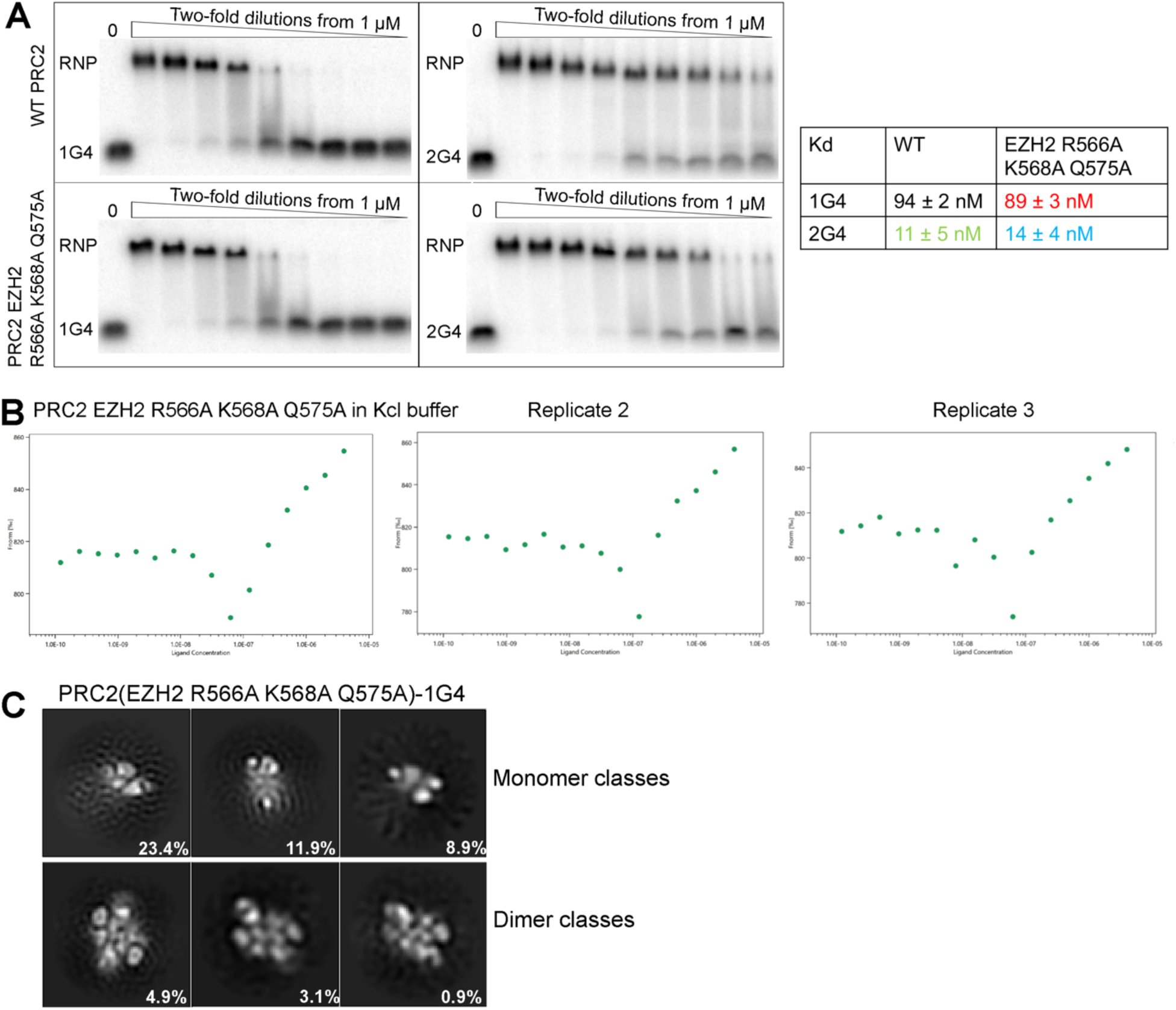
EZH2 R566A K568A Q575A does not impact 1G4-binding. (**A**) EMSA results of WT PRC2 and EZH2 R566A K568A Q575A binding to 1G4 and 2G4. (**B**) MST data of EZH2 R566A K568A Q575A binding 1G4 RNA. (**C**) 2D-class averages of EZH2 R566A K568A Q575A-1G4 RNA complex collected from streptavidin-affinity grid. Fraction of each class is highlighted at the bottom-right.

**Fig. S8.**
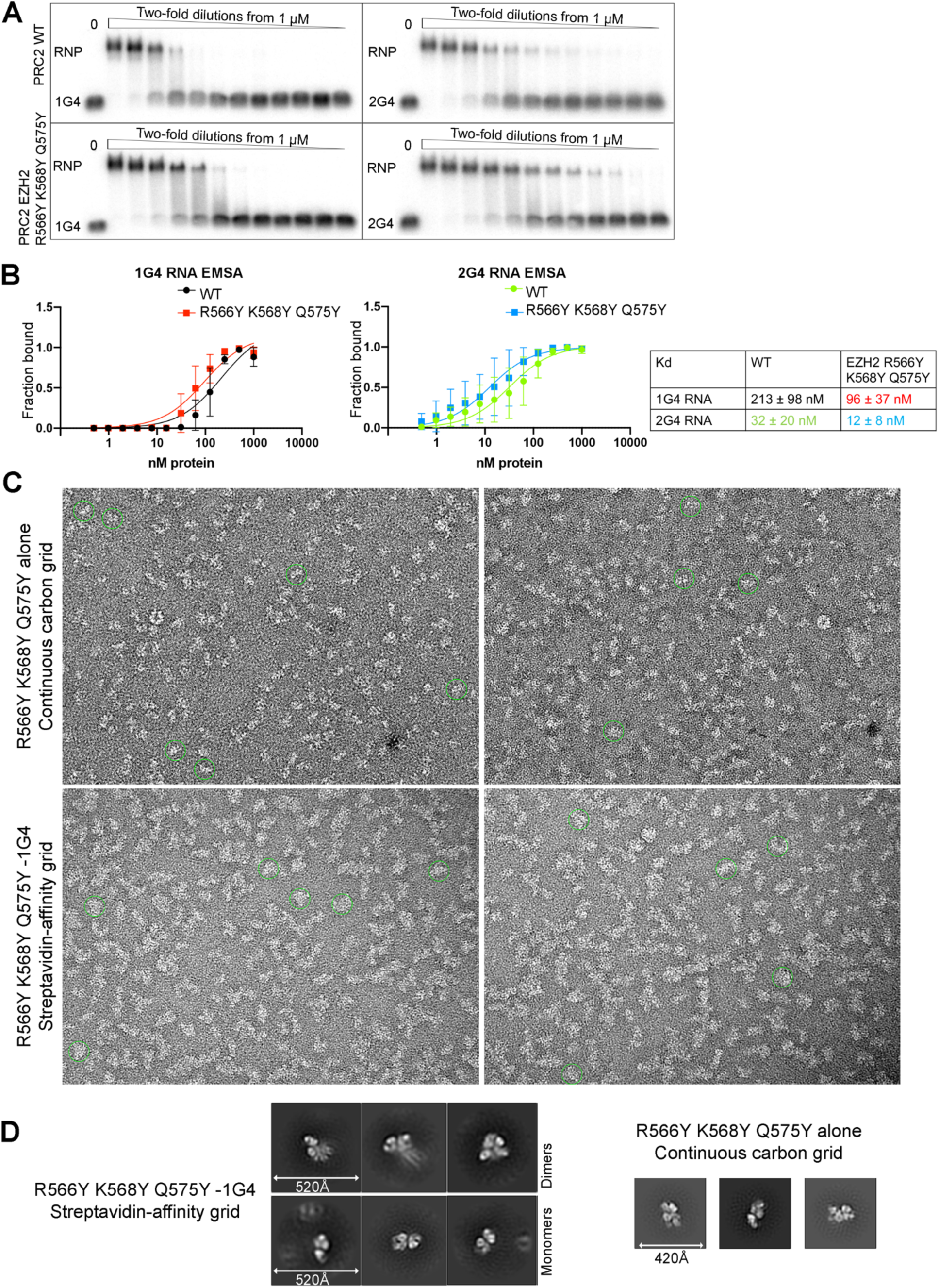
EZH2 R566Y K568Y Q575Y mutant of PRC2 has an increased RNA binding affinity and protein dimerization. (**A**) EMSA measurements of RNA binding to WT PRC2 and EZH2 R566Y K568Y W575Y. (**B**) Quantification of three EMSA replicates. (**C**) Negative staining EM images of EZH2 R566Y K568Y Q575Y on continuous carbon grid and 1G4-bound RNP on streptavidin-affinity grid after lattice-subtraction. Green circles have a diameter of 250Å to highlight size-difference between monomers and dimers. (**D**) Examples of 2D-class averages from EZH2 R566Y K568Y Q575Y-1G4 and mutant protein alone. Because this mutant formed very flexible dimers upon 1G4 binding, indicated by a smeared second protomer, we could not unambiguously group dimers by the regular 2D classification algorithm. Therefore, particles from raw images were manually counted to estimate the fraction of dimers.

**Fig. S9.**
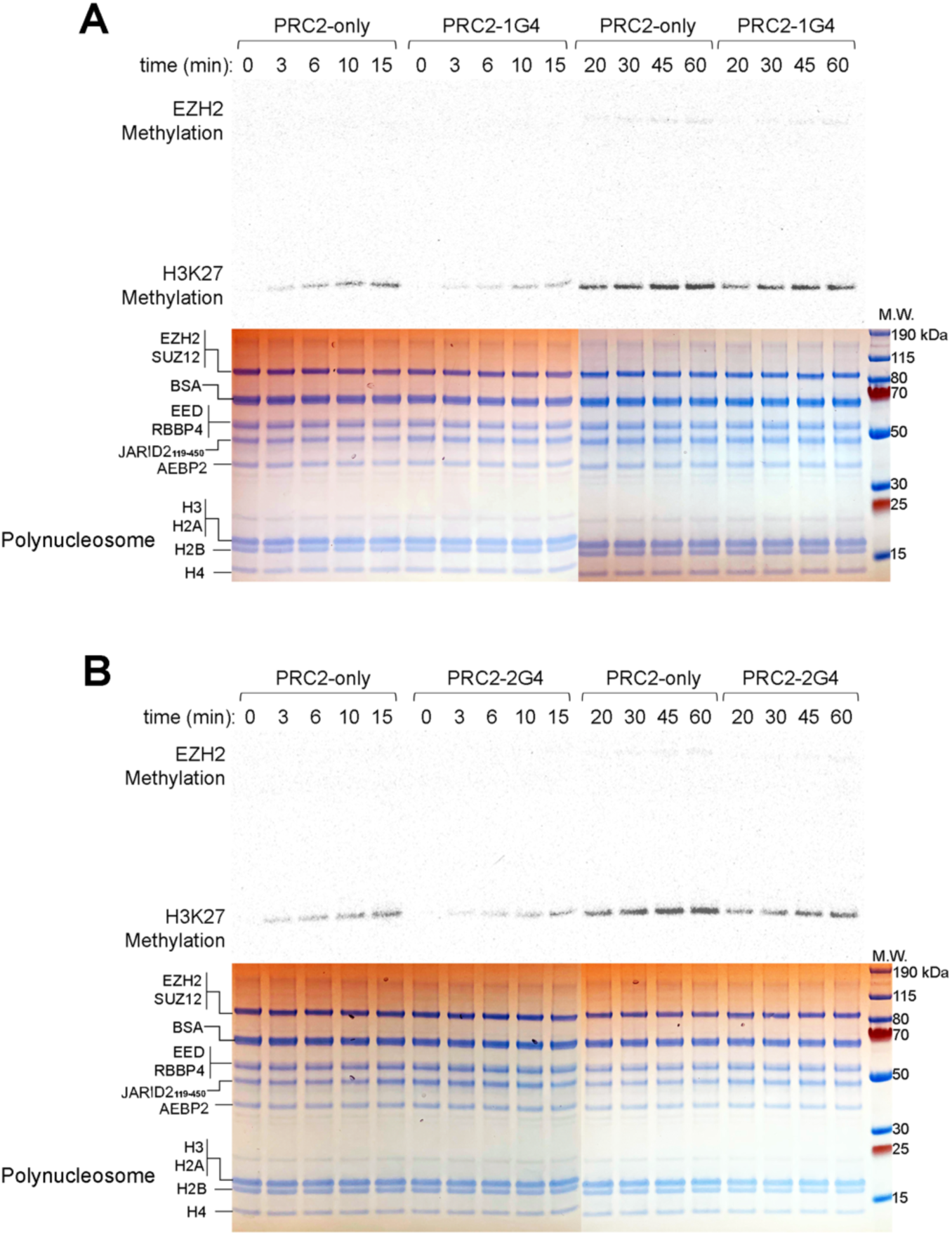
Methyltransferase activity assay of six-subunit PRC2 with polynucleosomes as substrates. We incubated PRC2, polynucleosomes, radiolabeled S-adenosyl methionine (SAM), and corresponding RNA or mock for different reaction times. Signal intensities were quantified with replicates and plotted in (Fig. 3F). Bottom images, Coomassie-staining of the same gels showing equal loading of each sample.

**Fig. S10.**
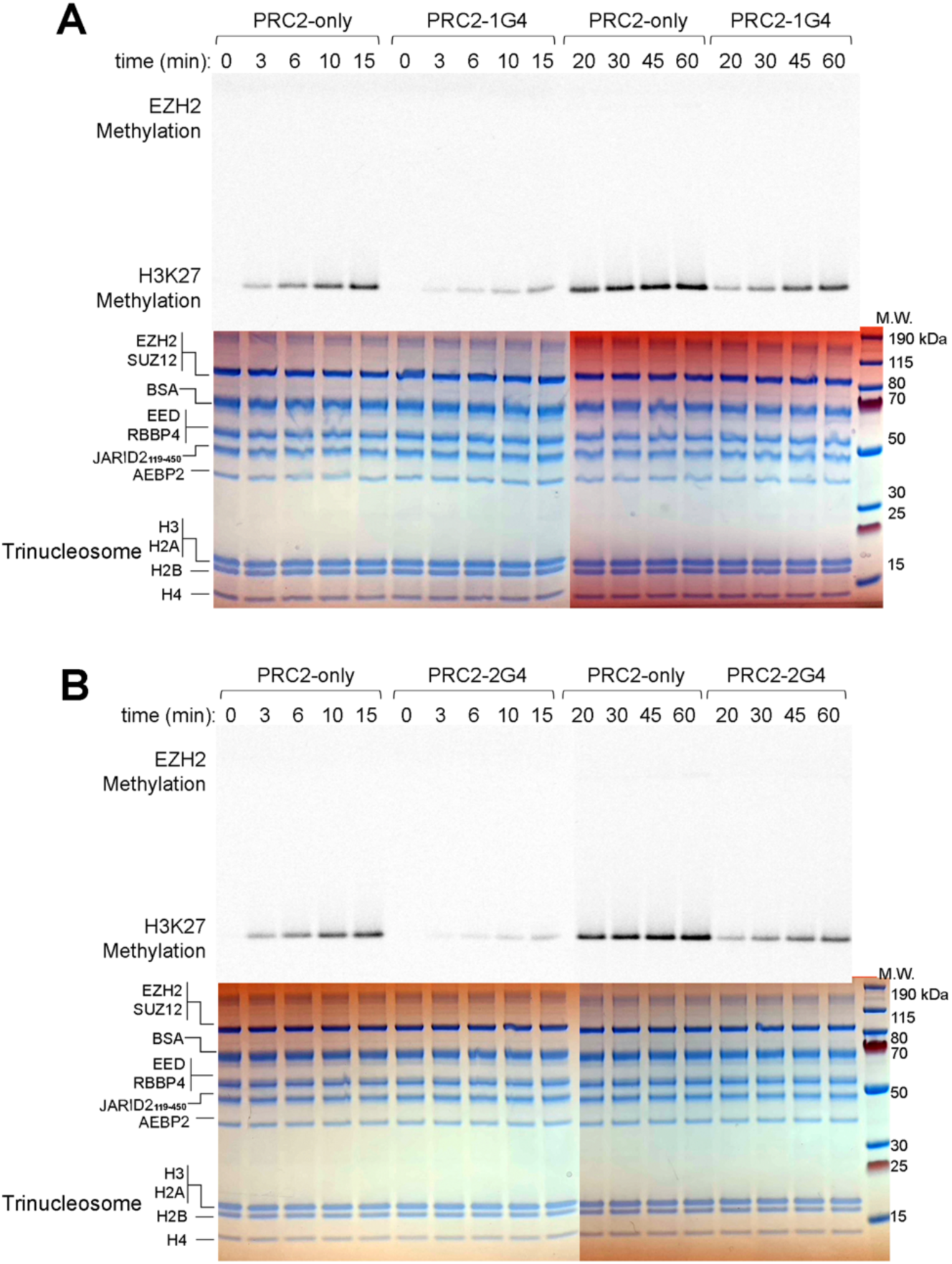
Methyltransferase activity assay of six-subunit PRC2 with trinucleosomes as substrates. We incubated PRC2, trinucleosomes, radiolabeled SAM, and corresponding RNA or mock for different reaction times. Signal intensities were quantified with replicates and plotted in (Fig. 3F). Bottom images, Coomassie-staining of the same gels showing equal loading of each sample.

**Fig. S11.**
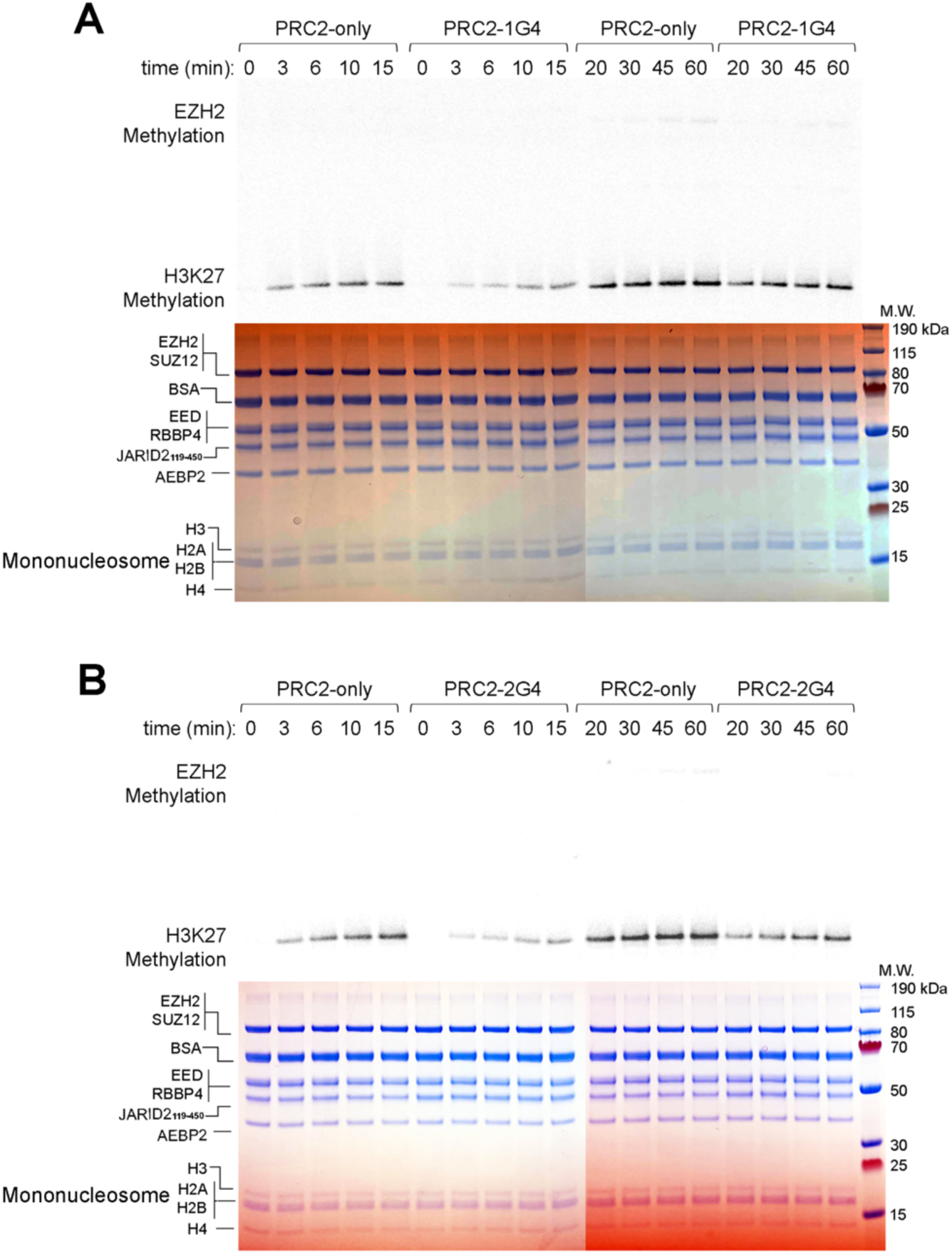
Methyltransferase activity assay of six-subunit PRC2 with mononucleosomes as substrates. We incubated PRC2, mononucleosomes, radiolabeled SAM, and corresponding RNA or mock for different reaction times. Signal intensities were quantified with replicates and plotted in (Fig. 3F). Bottom images, Coomassie-staining of the same gels showing equal loading of each sample.

**Fig. S12.**
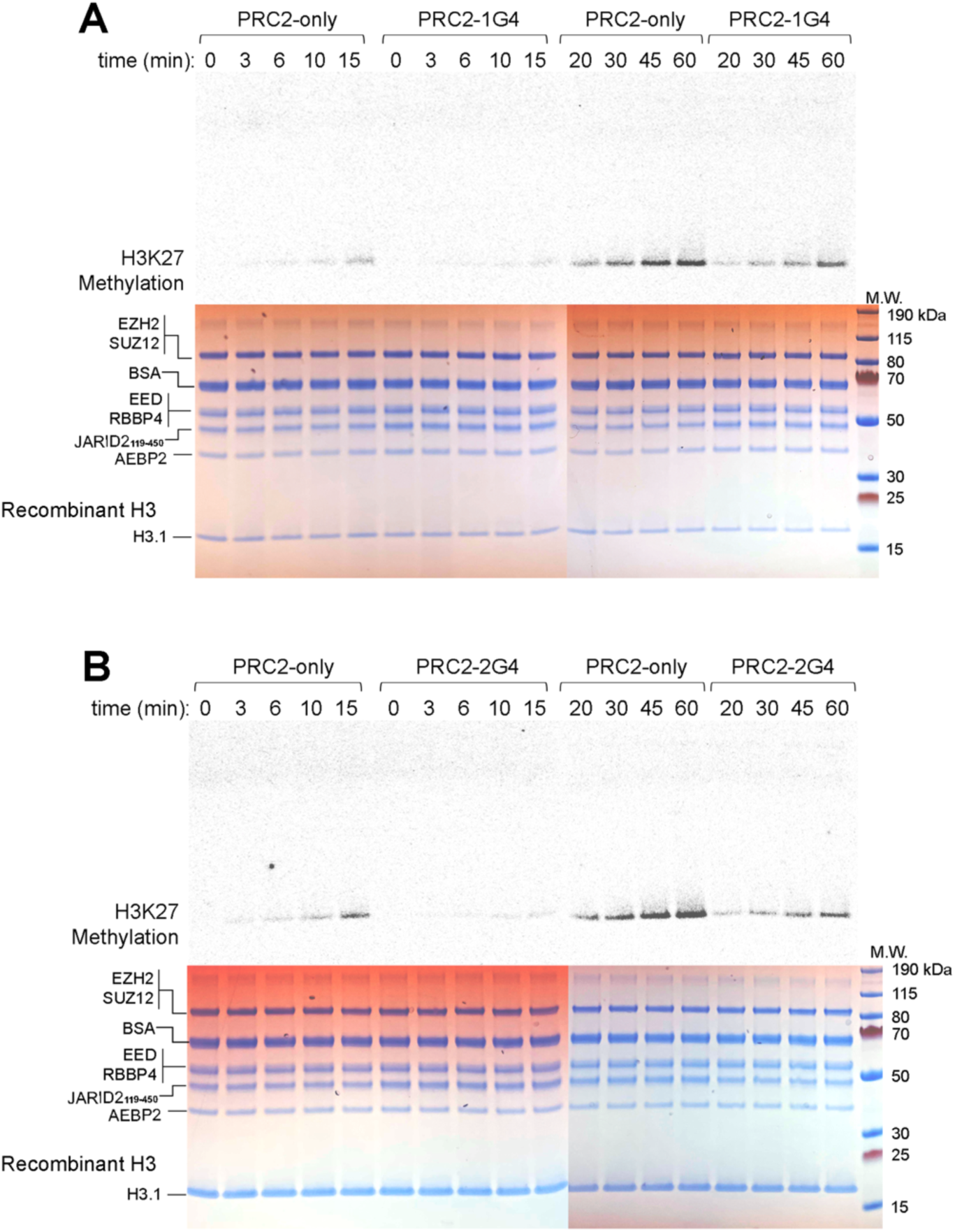
Methyltransferase activity assay of six-subunit PRC2 with recombinant H3 as substrate. We incubated PRC2, recombinant H3.1, radiolabeled SAM, and corresponding RNA or mock for different reaction times. Signal intensities were quantified with replicates and plotted in (Fig. 3F). Bottom images, Coomassie-staining of the same gels showing equal loading of each sample.

**Fig. S13.**
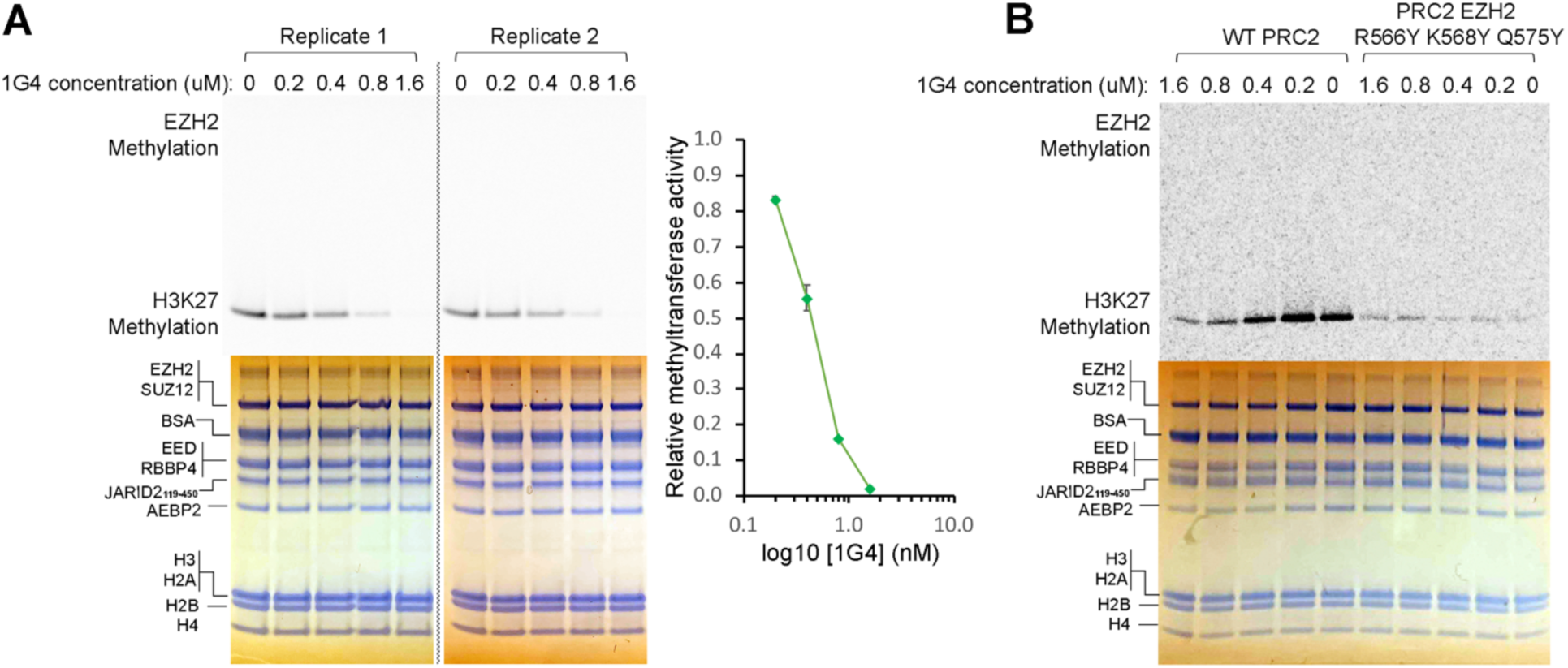
Inhibition of histone methyltransferase activity by 1G4 RNA. (**A**) Methyltransferase activity assays of PRC2 with serial dilutions of 1G4. Signal intensities were quantified and plotted on the right. Error bars are range of the values (N=2). (**B**) Methyltransferase activity assay to compare WT PRC2 with EZH2 R566Y K568Y Q575Y.

**Fig. S14.**
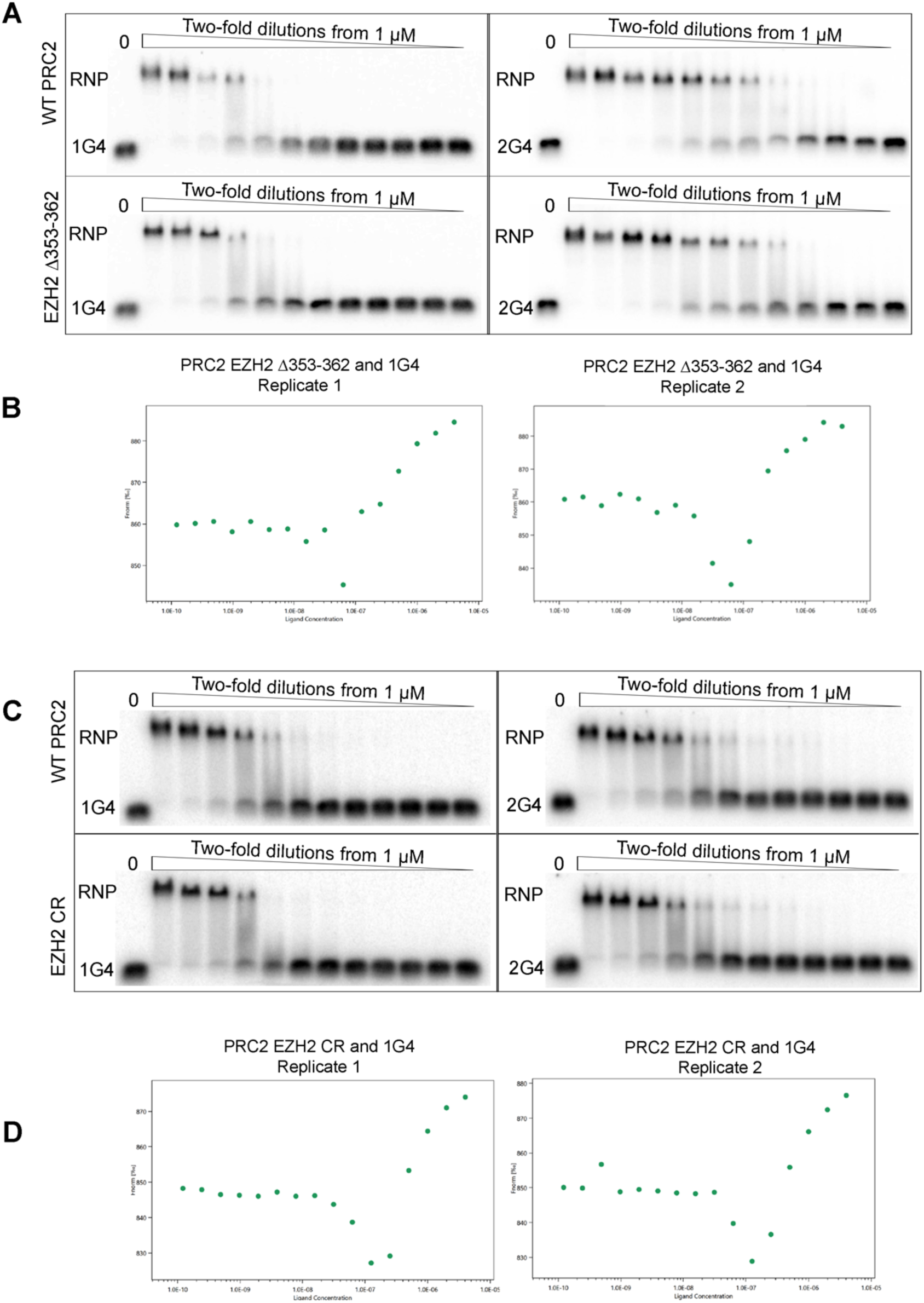
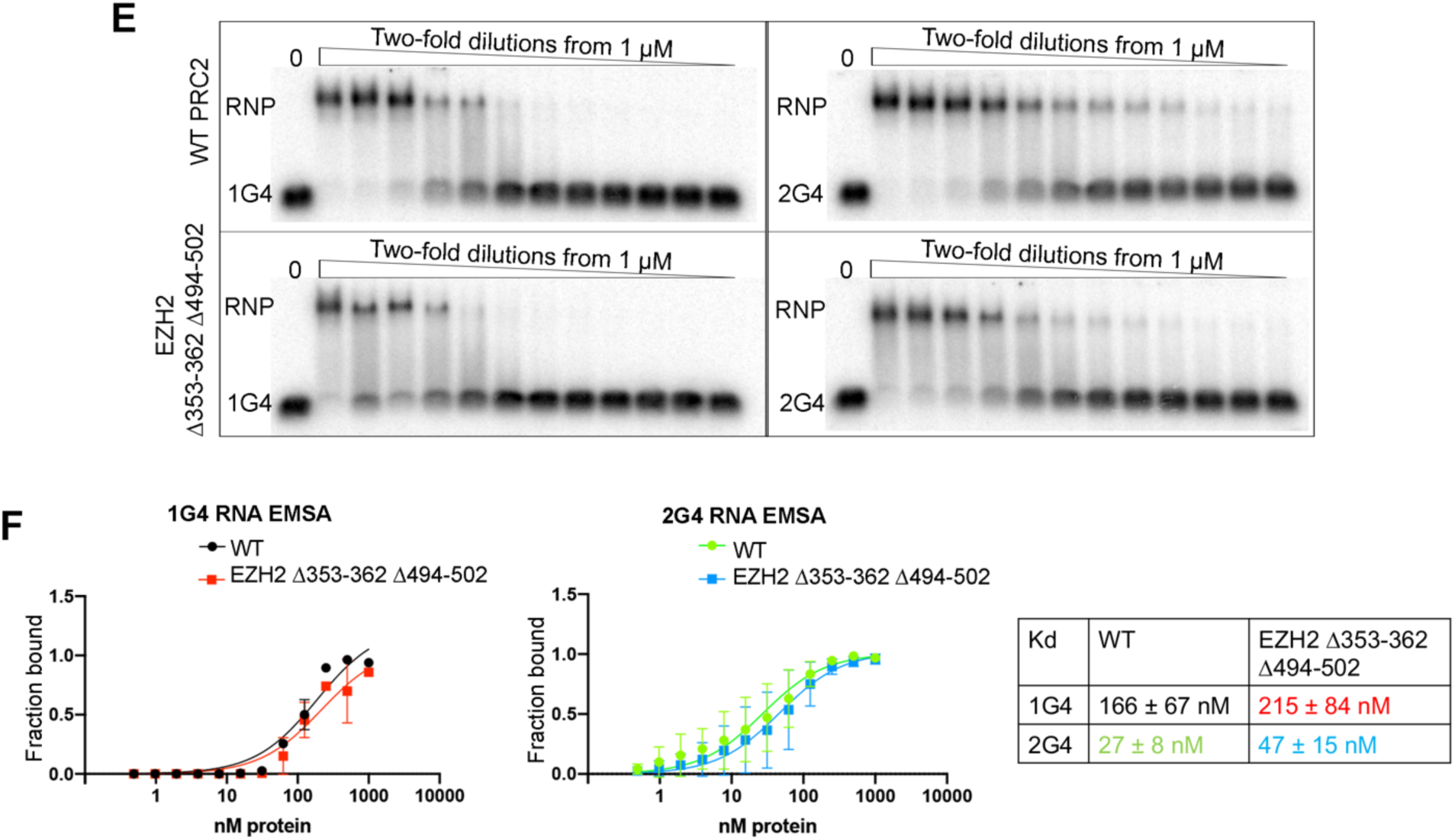
Characterization of EZH2 353-362 site in RNA binding. (**A**) EMSA results of WT and EZH2 Δ353-362 with 1G4 and 2G4 RNA. (**B**) MST data showing a two-stage curve for EZH2 Δ353-362-1G4 interaction. (**C**) EMSA results of WT and EZH2 CR mutant. (**D**) MST data of EZH2 CR-1G4 interaction. (**E**) EMSA results of WT and EZH2 Δ353-362 Δ494-502. (**F**) Quantifications of three EMSA replicates.

**Fig. S15.**
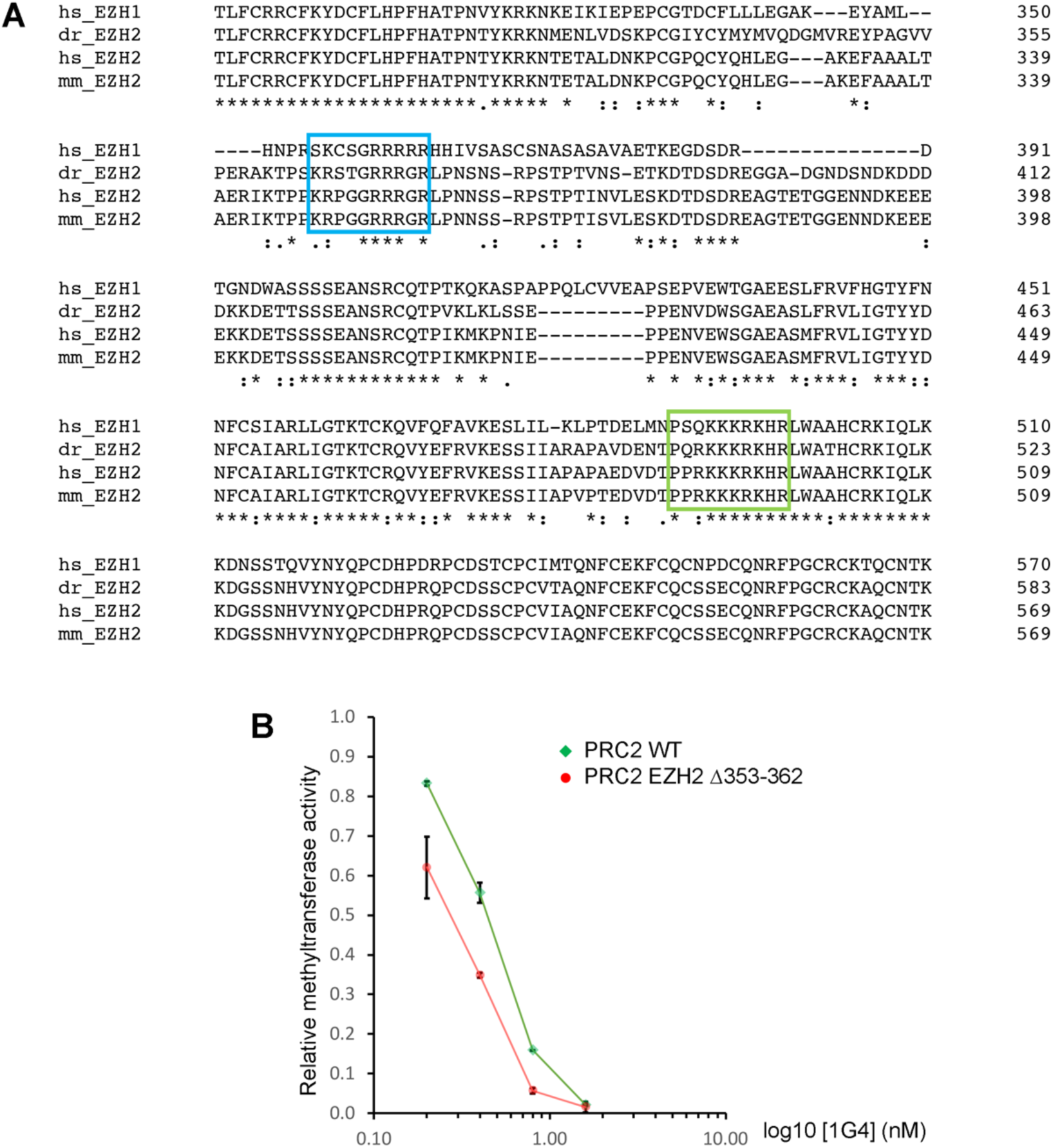
Characterizations of EZH2 353-362 site in nucleosome displacement. (**A**) Sequence alignment of EZH2 and EZH1. EZH2 353-362 and 494-502 sites are highlighted with blue and green circles correspondingly. *hs* is *Homo sapiens*; *dr* is *Danio rerio*; *mm* is *Mus musculus*. (**B**) Quantified methyltransferase activity assay from WT PRC2 and EZH2 Δ353-362 pre-incubated with various concentrations of 1G4 RNA. Error bars are range of the values (N=2).

**Fig. S16.**
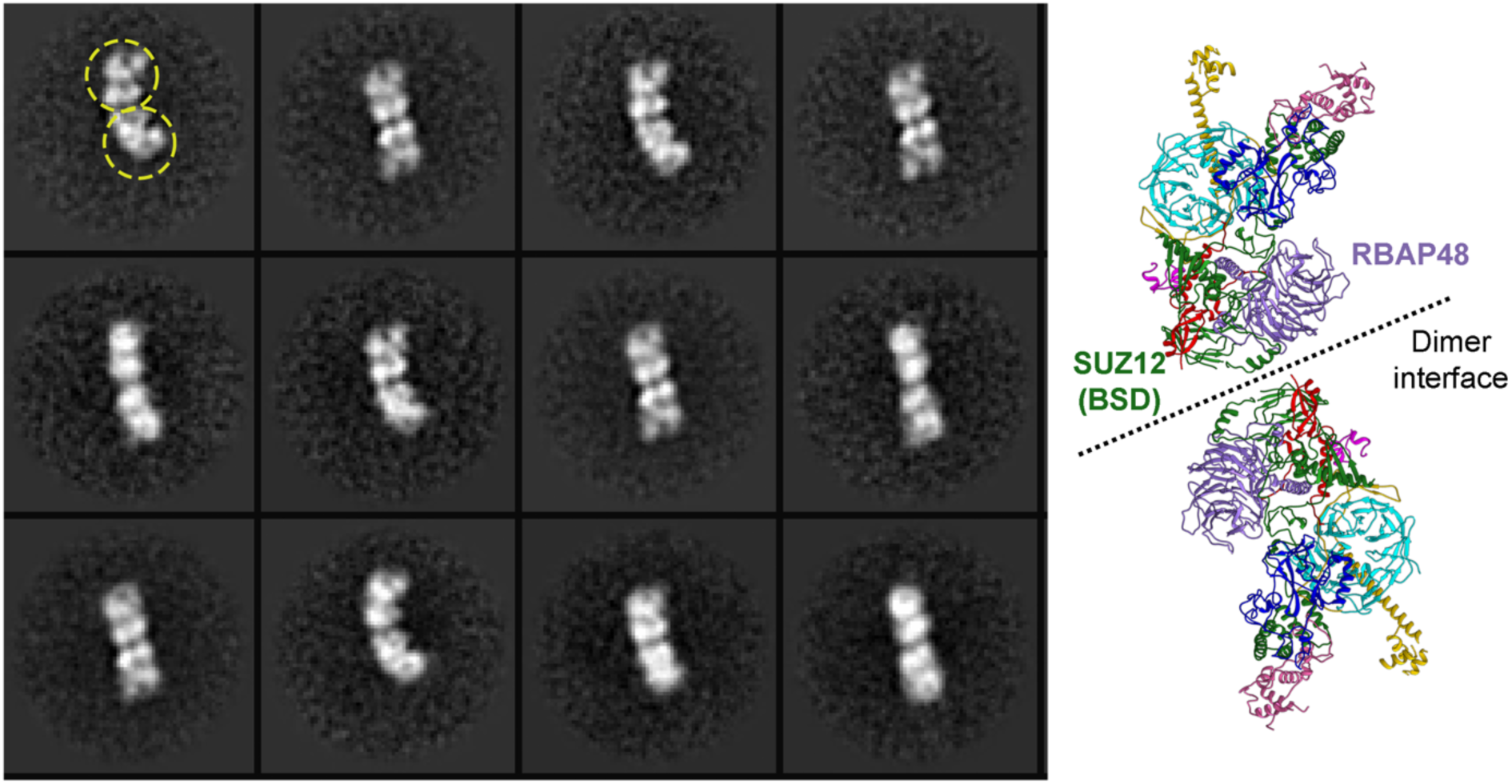
Dimerization of unbound six-subunit PRC2.2 has different dimer arrangement. 2D class averages of PRC2 dimers without RNA-binding exhibit a different dimer interface. Yellow circles highlight two PRC2 protomers. Dimerized PRC2 only occupied a small fraction (<10%) and was removed by gel filtration in protein purification.

**Table S1.** Cryo-EM data collection, refinement and validation statistics.

|  |  |  |  |
| --- | --- | --- | --- |
| **Data Collection** |  |  |  |
| Microscope | Krios |  |  |
| Voltage (kV) | 300 |  |  |
| Detector | K3 |  |  |
| Pixel Size (Å) | 0.422 |  |  |
| Defocus Range (µm) | -0.6 to -2.0 |  |  |
| Movies | 18,632 |  |  |
| Exposure Rate (e<sup>-</sup>/Å<sup>2</sup>) | 60 |  |  |
| **Reconstruction** | Body 1 | Body 2 | Consensus |
| Software | RELION 4 | RELION 4 | RELION 4 |
| Particles (final) | 217,196 | 217,196 | 217,196 |
| Accuracy Rotations (°) | 1.1 |  | 1.1 |
| Accuracy Translations (Å) | 0.55 |  | 0.72 |
| Map Resolution | 3.3 | 3.3 | 3.4 |
| Map Sharpening B-factor | -40 | -40 | -40 |
| **Coordinate Refinement** |  |  |  |
| Software | PHENIX | PHENIX | PHENIX |
| Map CC | 0.8 | 0.8 | 0.72 |
| Resolution Cutoff (Å) | 4 | 4 | 4 |
| Box Size (Å) | 147.82x110.6x129.74 | 156.33x111.66x106.34 | 171.21x203.12x146.75 |
| FSC<sub>model-vs-map</sub>=0.5 (Å) | 3.4 | 3.4 | 3.9 |
| **Model** |  |  |  |
| Number of Atoms | 12617 | 12617 | 25836 |
| Protein | 1846 | 1846 | 3692 |
| Nucleic Acid | 0 | 0 | 18 |
| **R.m.s deviations** |  |  |  |
| Bond Length (Å) | 0.002 | 0.002 | 0.009 |
| Bond Angles (°) | 0.495 | 0.459 | 0.628 |
| **Validation** |  |  |  |
| Molprobity score | 1.74 | 1.73 | 1.99 |
| Clashscore | 5.74 | 5.21 | 10.75 |
| Rotamer outliers (%) | 0.31 | 0.1 | 0.05 |
| Cβ outliers (%) | 0 | 0 | 0.03 |
| **Ramachandran** |  |  |  |
| Favored (%) | 93.38 | 92.99 | 93.16 |
| Allowed (%) | 6.40 | 6.73 | 6.62 |
| Outliers (%) | 0.22 | 0.28 | 0.22 |
| CaBLAM outliers (%) | 4.97 | 4.97 | 4.97 |

